# Symbolic regression enables coarse-grained model discovery of intracellular signalling dynamics

**DOI:** 10.64898/2026.08.20.745973

**Authors:** Theodore de Pomereu, Fabian Fröhlich

## Abstract

Cells respond to their environment through protein networks often dysregulated in cancer, making dynamical modelling crucial. Limitations in experimental data and computational resources motivate coarse-graining methods to build low-dimensional descriptions. Yet classical approaches to coarse-grained modelling rely on strong assumptions, leaving it unclear when partial experimental observations support reduced descriptions of system dynamics. Here we show that symbolic regression (SR) provides a data-driven way to test whether, and how compactly, the dynamics of a signalling system coarse-grain over the measured variables, and, when they do, infers mechanistically interpretable models. In synthetic enzyme systems, SR recovers Michaelis–Menten kinetics for the two-step mechanism and under three-step extensions. As data quality is degraded, SR simplifies toward effective kinetic laws while preserving correct theoretical limits. Applied to published time-resolved ERK phosphorylation data, SR identifies compact phospho-ERK rate laws in selected cancer-relevant gene overexpression contexts, yielding interpretable kinetic effects. A sparse neural ODE baseline requires few inputs where SR succeeds, but on average more where it fails, indicating that, where a reduced model is learnable at all, SR failure is associated with more complex dynamics that a simple mathematical model cannot describe. Together, these findings establish symbolic regression as a way to test when a compact coarse-grained description is warranted, generating hypotheses where one holds and motivating potential new measurements where it does not.

**Significance statement:** Cells process information through complex biochemical networks whose governing equations are usually unknown. Automated equation discovery has succeeded in physics and well-sampled biology, but whether it extends to sparse, noisy, partial data typical of these networks remains unclear. Here, from single-cell measurements of a growth-signalling pathway across dozens of cancer-relevant perturbations, we show it recovers compact models in many of them, some reproducing known regulatory biology and others proposing novel, testable hypotheses. A more expressive deep-learning baseline succeeds more often, but equation discovery’s failures cluster where that baseline uses more measured variables than a compact law can hold. Reducibility to a compact law becomes a measurable property, not a modelling assumption, indicating when dynamics compress and when they do not.

## Introduction

All models of biology are coarse-grained. At any level of description, further mechanistic detail could in principle be added, reflecting the effectively unbounded complexity of molecular systems^1^. Modelling therefore begins with two choices—a system boundary separating internal from external, and a scale of representation that resolves selected mechanisms while omitting finer spatial, temporal or organisational structure. Incomplete, noisy measurements and the computational cost of fitting and simulating large models make these choices unavoidable. The reduction is not merely pragmatic but conceptually grounded: because biological organisation is multiscale, appropriately coarse descriptions can capture effective interactions while remaining interpretable^2,3^, and can even predict better than fine-grained ones^4^. Yet a self-contained description at the level of the measured variables is not guaranteed, whether because of missing information or structural non-reducibility^5,6^.

This matters especially for intracellular signalling networks—systems of interacting proteins linked by phosphorylation and other post-translational modifications that drive cellular responses to the environment—whose high-dimensional, densely interconnected dynamics are hard to isolate from their intracellular context^7^, and where local molecular detail can reshape system-level behaviour^8^. Cancer illustrates this point: single molecular alterations in key regulatory proteins can subvert network logic, rewiring growth-control decisions to permit uncontrolled proliferation^9,10^. Conversely, the efficacy of drugs targeting those alterations depends on how the network has been rewired, not on the target alone^11^. Robust dynamical models linking perturbations to network-level outcomes are therefore clinically relevant.

The dominant paradigm uses mechanistic models in which measured species are linked by ordinary differential equations (ODEs) derived from prior knowledge, typically with mass-action kinetics; parameters are fit and the equations integrated to predict time courses^12^. Its appeal is generality: once the ODE is fixed, it can be simulated across many perturbations and out-of-distribution contexts. In practice, resolution and precision are limited by restricted observability, parameter non-identifiability^13^, uncertain prior knowledge^6^, sensitivity to cellular context^7,11^, and rate-law assumptions that are often difficult to justify in vivo^14–16^.

These constraints have long motivated principled coarse-graining of biochemical reaction networks. Much of this work exploits time-scale separation, from classical quasi-steady-state reductions^17,18^ (e.g. Michaelis-Menten kinetics) to frameworks clarifying when eliminations are valid and how the reduced dynamics depend on what is treated as fast versus slow^19^. Other methods include lumping^20,21^, sensitivity- or optimisation-based pruning^22–24^, profile-likelihood reduction^25^, manifold boundary approximation^26^, and projection-based reductions from singular value decomposition (SVD) methods such as balanced truncation^27^ to operator-theoretic Koopman embeddings^28^—reflecting that the “right” reduction depends on the modelling goal and the available observables. Most of this literature, however, treats reduction as post-hoc simplification of a detailed mechanistic model under idealised assumptions, yielding approximations tailored to particular observables or regimes^29,30^. These approaches do not address cases in which a detailed mechanistic has yet to be developed, whether because doing so is computationally or conceptually challenging. We therefore focus on as an integral part of the model-construction process.

Scientific machine learning (SciML) offers tools for model discovery that learn such descriptions directly from data, informing not only parameter values but the functional form of the dynamics: black-box models such as neural ODEs^31^, physics- or biology-informed neural networks^32,33^, and hybrids such as universal differential equations that add learned residuals to mechanistic components^34^. Symbolic regression (SR) offers a particularly explicit route, searching directly for compact analytical expressions^35^ through strategies including genetic-programming expression-tree search (PySR^36^), physics-guided reconstruction (AI-Feynman^37^), reinforcement-learning–guided program search (Deep Symbolic Optimisation^38^; DSO), and neural-symbolic hybrids such as Kolmogorov-Arnold Networks^39^ (KANs). Across physics, chemistry and engineering, SR has recovered both known laws from classic systems^37,40,41^ and novel models from experimental data^42,43^. Though usually introduced for static input–output relationships, it extends to dynamics by discovering parsimonious differential equations from time-resolved trajectories^40,44,45^. In biology, where series are noisy and partially observed, SR has yielded compact phenomenological descriptions, but mainly for well-resolved time series such as bacterial growth and fermentation^46,47^ rather than sparse molecular interaction data.

Here we ask whether SR can identify coarse-grained models of signalling from measured variables and, more broadly, whether SR’s success or failure can serve as a data-driven readout of how coarse-grainable those dynamics are; when it cannot, whether that failure is itself informative; and when it can, what the models reveal about feedback, control and pathway coupling. We first benchmark state-of-the-art SR on synthetic enzyme-kinetics trajectories, showing that it rediscovers Michaelis-Menten rate laws and remains robust to mechanistic extensions and to realistic degradations in noise level, sampling, kinetic-regime coverage and choice of observable. We then apply the same framework to time-resolved perturbational phospho-proteomic data, inferring coarse-grained ODEs for ERK phosphorylation across cancer-relevant overexpression contexts. Compact, predictive rate laws emerge for a subset of contexts; where SR fails, a sparse neural ODE requires more inputs to fit the same data, indicating dynamics that on average resist compression into a compact model rather than a limitation of SR. Where SR succeeds, the learned rate laws yield mechanistic hypotheses about feedback and pathway coupling.

## Results

### Symbolic regression identifies and extends existing coarse-grained models from classical enzyme kinetics

We first asked whether SR could recover established coarse-grained models from data, and where such recovery fails. As a test case, we investigated whether SR could rediscover the Michaelis-Menten rate law from synthetic time-resolved measurements generated by mechanistic enzyme models. Michaelis-Menten gives a classical coarse-grained description of enzymatic catalysis^48^:

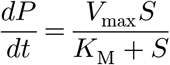

where *S* is free substrate, *P* product, *E*_tot_ total enzyme, *k*_cat_ the catalytic rate constant, *k*_off_ the enzyme-substrate unbinding rate, *K*_D_ the dissociation constant, *V*_max_ = *k*_cat_*E*_tot_ the reaction rate and *K_M_* = (*k*_cat_ + *k*_off_ )/*K_D_k*_off_ the Michaelis constant (Fig. 1A). We compared four SR methods (AI-Feynman^37^, PySR^36^, DSO^38^, KANs^39^) on synthetic data from a closed-system two-step mechanism—no inflow or outflow, dynamics driven solely by the reaction—comprising reversible enzyme-substrate binding followed by irreversible catalytic product formation (Fig. 1A), which admits the Michaelis-Menten approximation under the quasi-steady-state assumption^17^. Parameters and initial conditions were drawn log-uniformly over 10^-5^ to 10^3^ and trajectories recorded at steady state; each method took *S*, *E*_tot_, *k*_cat_, *k*_off_ and *K_D_* as inputs and predicted the renormalised product rate *dP* /*dt* (Methods). Accuracy was scored by median relative absolute error (MdRAE), chosen for robustness to outliers. PySR (yellow) achieved the lowest error (MdRAE = 1.50×10^-9^%; Fig. 1B, Table 1) and recovered the exact Michaelis-Menten equation, whereas AI-Feynman (red), DSO (green) and KAN (black) remained far less accurate (median relative absolute error 100.00%, 756.38% and 92.24%, respectively) (Fig. 1B). We attribute this to PySR’s custom log-space loss, suited to targets spanning orders of magnitude^36^: the others required log-transforming the target and adding a log operator, which yields many invalid candidates when intermediate values turn non-positive, hindering the search. SR implemented in PySR therefore recovers a canonical coarse-grained biochemical model directly from data.

**Figure 1.**
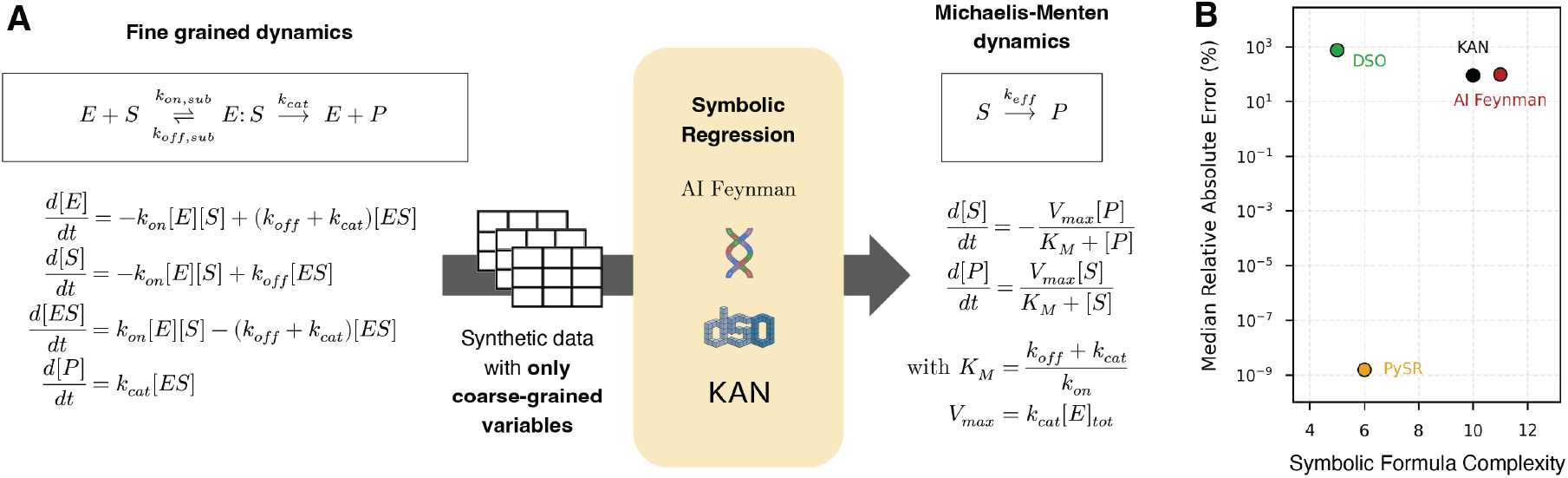
Symbolic regression recovers the Michaelis-Menten equation at low complexity and low error, with performance varying sharply across methods. (**A**) The reduction task. Left, the two-step enzyme-catalysis mechanism and its mass-action description; right, the target coarse-grained Michaelis-Menten form. (**B**) Each point is the best expression recovered by one of four symbolic-regression (SR) methods in a single run (seed 42), positioned by symbolic complexity (x) and test-set median relative absolute error (MdRAE, y); lower is better on both axes. Training-set sizes differ by method following method-specific practice (PySR n = 3,000; AI-Feynman n = 5,000; DSO n = 38,400; KAN n = 4,000), each under a fixed 10,800 s (3 h) budget. PySR recovers the Michaelis-Menten form, whereas AI-Feynman, DSO and KAN — run under their standard log-transformed-target settings — remain high-error.

**Table 1.**
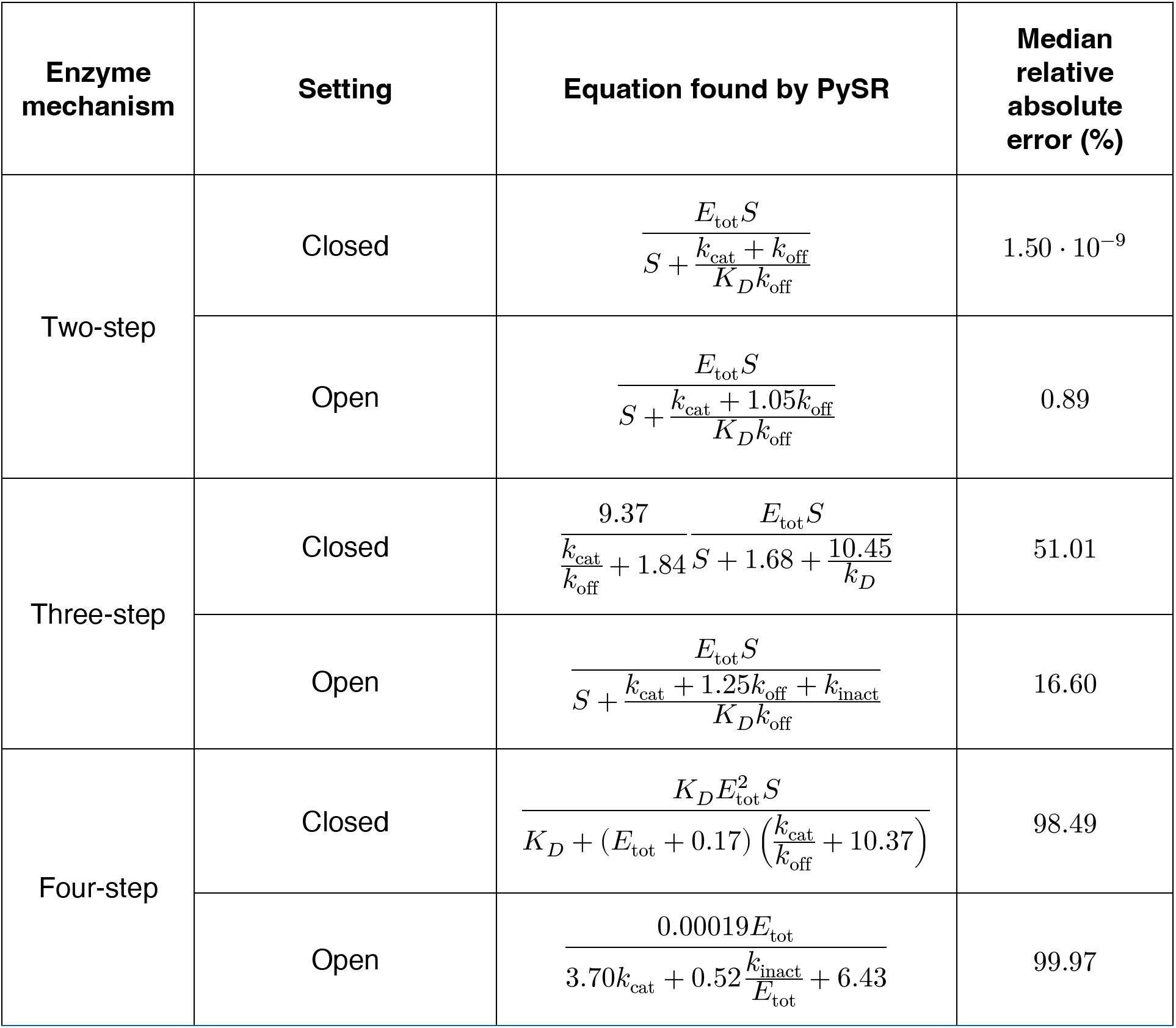
Equations recovered by PySR for synthetic enzyme mechanisms in closed and open systems, with test-set median relative absolute error.

To assess robustness beyond the minimal mechanism, we evaluated PySR—the top performer above—on more complex schemes adding reversible enzyme–product binding and enzyme dimerisation, a common feature of kinase activation in signalling^49^ (Fig. 2A). For the three-step mechanism, PySR recovered a coefficient-1 Hill function preserving the characteristic substrate dependence, with renormalising factors absorbing the extra steps (Table 1, Fig. 2D), but a substantially higher MdRAE (51.01%). For the four-step mechanism it converged to a high-error (98.49%), high-complexity expression (21, versus 14 for Michaelis-Menten; Methods) (Table 1, Fig. 2C). Because no gold-standard coarse-grained description exists for these mechanisms, it remains unclear whether this reflects a limitation of SR or an inherent lack of coarse-grained structure. SR therefore recovers coarse-grained kinetic laws as complexity increases, but only up to a point.

**Figure 2.**
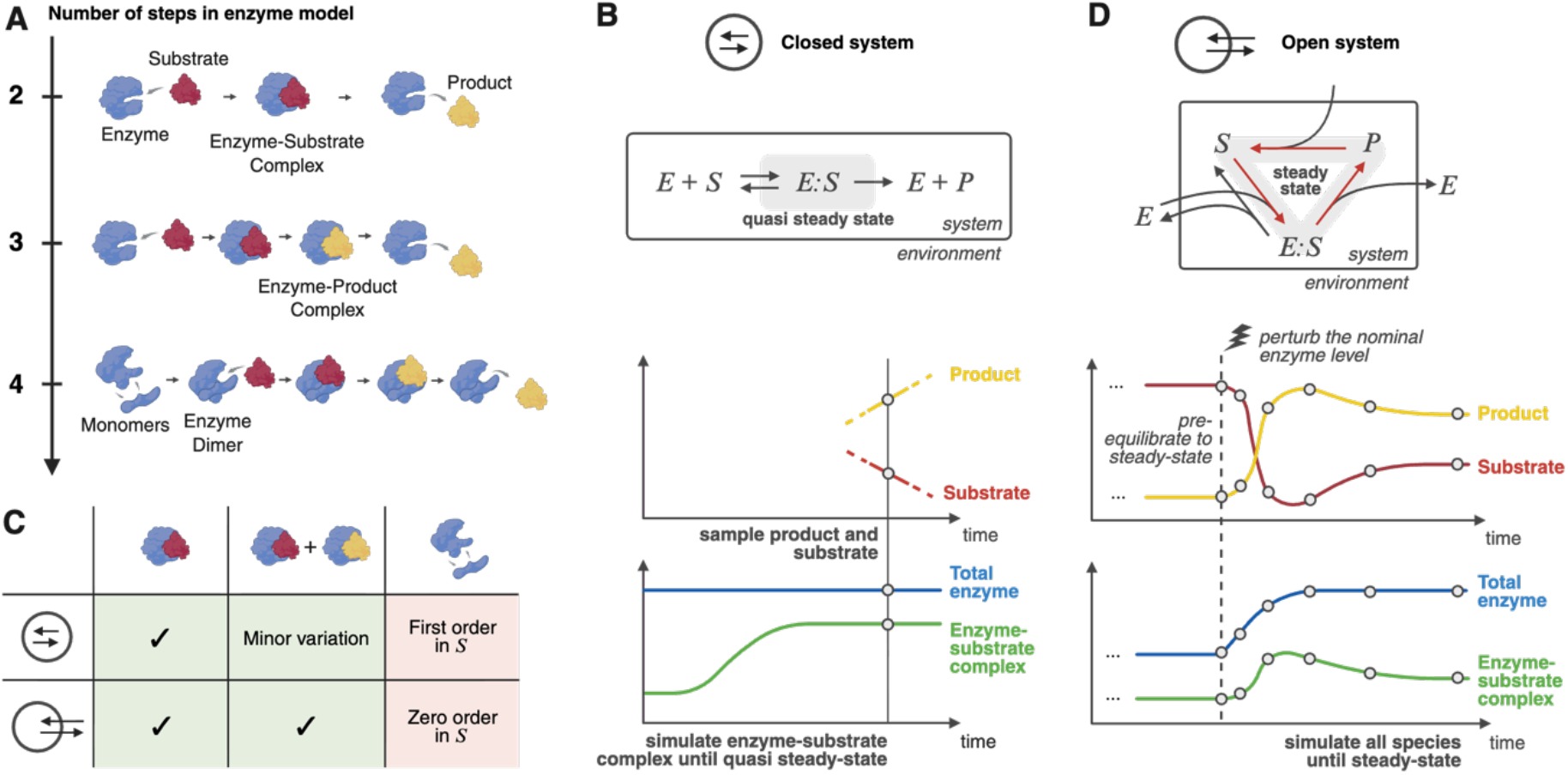
Synthetic enzyme systems used to evaluate symbolic regression across mechanistic complexity. **(A)** Three mechanisms of increasing complexity: two-step enzyme-substrate, three-step with reversible enzyme-product binding, four-step with enzyme dimerisation. (**B**) Closed-system setting. The black box marks the modelled system boundary; open circles at the vertical sampling line mark the state recorded for SR once the enzyme-substrate complex reaches quasi-steady state. (**C**) SR results by mechanism (columns) and setting (rows), one run per cell (seed 42). Check marks, exact recovery of the Michaelis-Menten form; text labels, the dominant alternative reduced behaviour recovered. (**D**) Open-system setting. Arrows crossing the boundary denote coupling to the environment; an initial simulation (ellipsis) runs to pre-equilibration, after which trajectories are sampled at the marked timepoints (open circles) until steady state.

We next asked whether SR adapts when Michaelis-Menten assumptions are violated. Relaxing the closed-system assumption, we extended each mechanism to an open-system setting in which enzyme, substrate and product concentrations are regulated by the environment (Fig. 2D), adding enzyme activation and inactivation at rates *k*_act_ and *k*_inact_ alongside degradation reactions coupling substrate and product turnover. These violate conservation of enzyme and substrate pools and timescale separation^50^. Data captured the full dynamics from an initial steady state to a new one following perturbation of *k*_act_ (Methods). For the three-step mechanism PySR recovered a form closer to Michaelis-Menten, with lower MdRAE (16.60%) than in the closed system, and still recovered Michaelis-Menten for the two-step mechanism at an increased but low MdRAE (0.89%) (Table 1, Fig. 2C). The four-step mechanism was again not amenable to coarse-graining (Table 1, Fig. 2C). SR therefore recovers classical enzyme kinetics under idealised conditions and adapts their functional form as mechanistic assumptions are relaxed.

### Symbolic regression preserves Michaelis-Menten kinetics and infers principled departures as measurement conditions degrade

We next asked how SR behaves when data quality and coverage are imperfect, as in most biological measurements. We degraded the synthetic dataset in five ways: (i) additive measurement noise, (ii) reduced training-set coverage, (iii) increased mismatch from Michaelis-Menten kinetics, (iv) alternative measured observables, and (v) restriction to specific kinetic limits. We chose the open-system three-step mechanism as a testbed as it was the most significant departure from classical assumptions under which SR still recovered the Michaelis–Menten equation (16.6% MdRAE). Each test used three seeds, so that variation among regimes reflected the degradation rather than initialisation. Throughout we benchmarked SR against two extremes: a rigid Michaelis-Menten “one-size-fits-all” solution and a flexible black-box multilayer perceptron (MLP).

We first asked how measurement quality affects the structure of recovered equations. The first two experiments added Gaussian noise (1%, 10%, 100% of the data standard deviation; Fig. 3A, purple boxes) and varied training-set size (10%, 50%, 100%, 200% of the standard 2,400-sample set; Methods; Fig. 3A, cyan boxes). Numerical loss scaled predictably with degradation (Fig. 3A); our interest was instead whether mechanistically meaningful functional forms survive. Both experiments show they do across a broad range of realistic data quality. At 1% noise SR recovered the exact Michaelis-Menten form in all three seeds. At 10% it returned Michaelis-Menten or near-identical variants, so the saturating substrate dependence remained identifiable. Only at 100% noise did SR collapse to a substrate-independent constant, consistent with substrate dependence being obscured by measurement variability. Dataset size behaved analogously: at 100% and 200% SR reliably returned the exact equation, whereas 10% and 50% increased inter-seed variability while largely preserving Michaelis-Menten-like structure (complexities 13-14; Fig. 3A, Tables S1 and S2). At 10%, one seed simplified to a linear substrate dependence (complexity 13; Table S2), reflecting reduced information content. Against the baselines SR was intermediate: at 0 and 1% noise and at 100% and 200% dataset size it closely matched the analytical model (Fig. 3A-B), while at higher noise or smaller datasets it fell between the analytical baseline and the MLP. Degradation is therefore controlled: SR preserves Michaelis-Menten structure under moderate loss of data quality and finds lower-complexity equations only when the data no longer support saturation.

**Figure 3.**
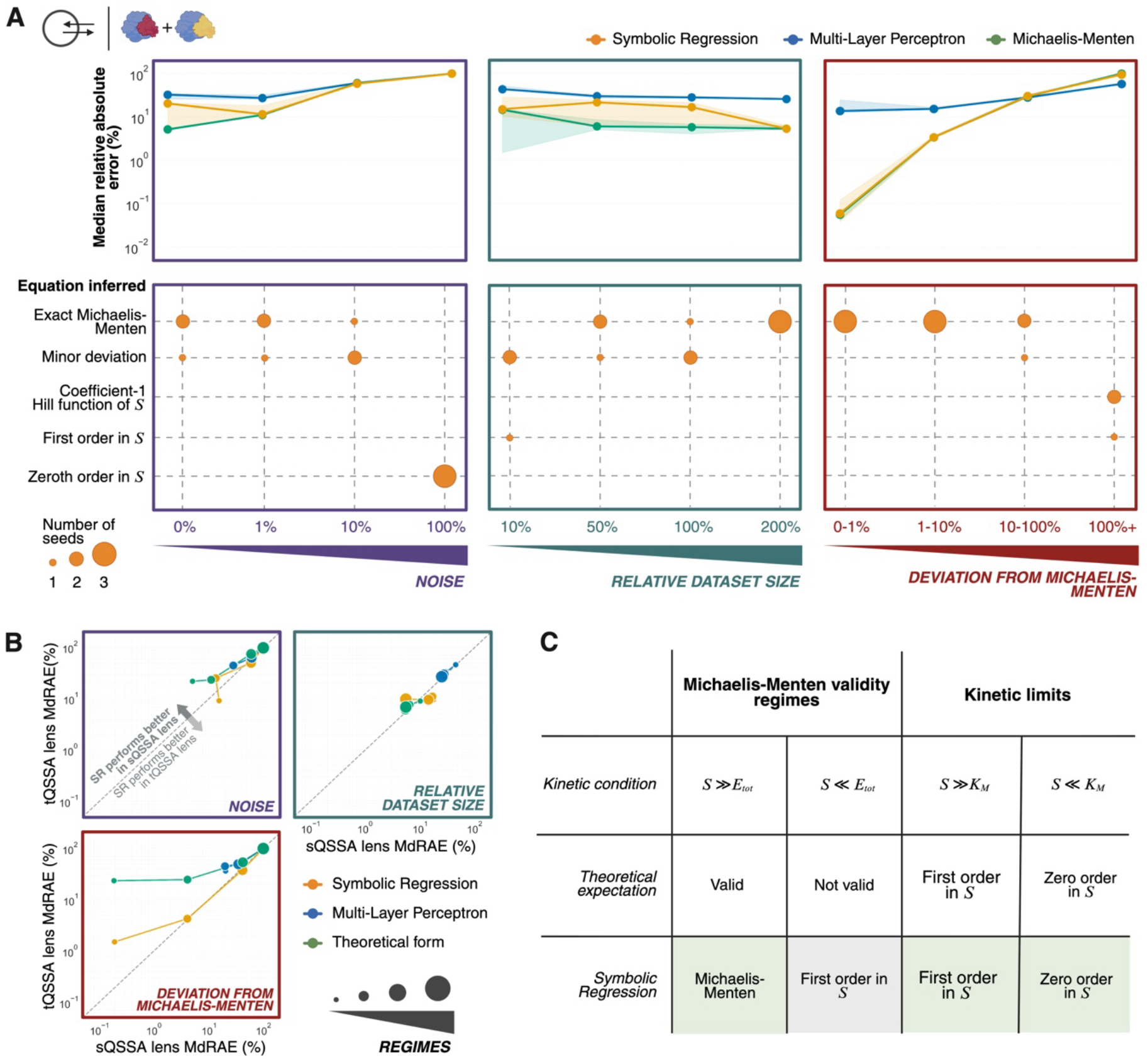
Symbolic regression performance across synthetic data degradation experiments. All panels use the open-system three-step mechanism with n = 3,000 samples per condition (2,400 training after the 20% test test split; the training-set-size experiment scales this base) and three seeds (42–44). (**A**) Three degradation experiments : measurement noise (left, purple), training-set size relative to the standard dataset (middle, cyan), and deviation of the data subset from the Michaelis-Menten regime (right, red). Upper row, seed-median test-set MdRAE for SR, the multilayer perceptron (MLP) baseline and the analytical Michaelis-Menten form; bands span the range across the three seeds. Lower row, bubble position gives the equation form found by SR and bubble size the number of seeds returning it (1-3). (**B**) Predictive error under the free-substrate (sQSSA) and total-substrate (tQSSA) lenses across the same regimes. Each point is one regime, point size tracks degradation level as in (*A*), and the dashed diagonal marks equal error. (**C**) Theoretical expectation versus recovered form. Left, validity of the Michaelis-Menten approximation against the substrate-to-enzyme ratio; right, the expected first- and zero-order limits against the substrate-to-Michaelis-constant ratio. Bottom row, the form SR inferred in each regime.

We next tested adaptation when the data are constitutively not Michaelis-Menten. The third experiment partitioned data by deviation from the Michaelis-Menten equation (Fig. 3A, red box), quantified as the absolute log error between the ground-truth rate and a Michaelis-Menten predictor computed directly from the features, into four regimes: low (< 0.01), medium (0.01–0.1), large (0.1–1.0) and very large (≥ 1.0). SR recovered Michaelis-Menten consistently in the first three, so moderate departures did not preclude recovery of the coefficient-1 Hill form (Fig. 3A). Only in the very large regime did expressions shift towards simpler coefficient-1 Hill approximations (complexity 9) and linear relationships (complexity 7; Table S3), consistent with the data being poorly described by a single Michaelis-Menten law. Across all regimes SR (yellow) matched Michaelis-Menten (green) in MdRAE, so more parsimonious forms cost no predictive power. SR strongly outperformed the MLP (blue) in the low and medium regimes and matched it in the large and very large (Fig. 3A). SR therefore recovers Michaelis-Menten only when the sampled regime supports it, and otherwise contracts to equally predictive lower-complexity expressions.

Our fourth experiment tested robustness to the choice of readout. We compared two measurement “lenses”: a free-substrate lens measuring only unbound substrate, and a more realistic total-substrate lens measuring free and enzyme-bound substrate together, since assays typically report total protein abundance rather than free monomer^51^. This alters the appropriate theoretical description: with total substrate *S_T_* = *S* + *ES* , the reduced kinetics follow not the standard Michaelis-Menten form under the standard quasi-steady-state assumption (sQSSA) but the total quasi-steady-state form^15^ (tQSSA):

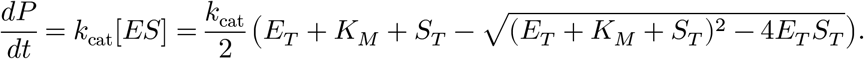

We crossed this lens shift with the noise, size and deviation experiments. Under size reduction the two lenses diverged and neither dominated (Fig. 3B, cyan box). Under noise and deviation, tQSSA performance was comparable to sQSSA (Fig. 3B, purple and red boxes), but SR never recovered the exact analytical tQSSA formula, favouring lower-complexity approximations that retained high accuracy (Fig. 3B, Tables S4-6). For the deviation experiment this follows from the setup, since data were stratified by closeness to the sQSSA form while the total-substrate representation obeys a different, algebraically more complex reduction. At low noise, where no such stratification applies, the same result indicates that SR generally favours parsimony over the expected tQSSA form. SR always matched or beat the MLP (blue, Fig. 3B). SR is therefore robust to the choice of measured observable, and prioritises functional parsimony over recovering complex theoretical derivations.

Finally, we subsetted the data to specific kinetic limits defined by enzyme-to-substrate (*E*/*S*) and substrate-to-Michaelis-constant (*S*/*K_M_* ) ratios (Fig. 3C). SR returned the expected limiting behaviours^52^: linear dependence in the first-order regime (*S*/*K_M_* ≪ 1) and an approximately constant rate in the zero-order regime (*S*/*K_M_* ≫ 1) (Fig. 3C). Where the quasi-steady-state assumption was violated (*E*/*S* ≫ 1), SR no longer inferred a coefficient-1 Hill form and favoured linear kinetics (complexities 9-13; Table S7). SR therefore finds forms matching the dominant kinetic behaviour in the data rather than defaulting to Michaelis-Menten or failing. Across all degradation experiments, SR identifies the coarse-grained kinetic law the data support, matching classical models where their assumptions remain informative and adapting to interpretable alternatives where they break down.

### Symbolic regression enables coarse-grained model discovery in experimental ERK signalling data across cancer-relevant perturbations

To test whether SR can extract interpretable kinetic relationships from real biological data, where the governing rate law is unknown, we turned to intracellular signalling. Using published time-resolved perturbational phospho-proteomic data from Lun *et al.*^51^ (Fig. 4A), we derived coarse-grained models of ERK phosphorylation—an enzyme-catalysed reaction embedded in a dense signalling network for which no universal mechanistic rate law exists^14^ (Fig. 4B). Lun *et al.* profiled 40 perturbation contexts^51^: 32 cancer-relevant kinase/phosphatase overexpression constructs and 8 controls. Cells were stratified along a GFP-tagged overexpression gradient and, after EGF addition at time 0, profiled by single-cell mass cytometry as cross-sectional snapshots at six timepoints over one hour, measuring a panel of 32 phospho-markers^51^. For each context, and across three seeds, we asked whether SR could infer parsimonious rate laws predicting ERK phosphorylation from seven local signalling inputs (p-ERK, p-MEK, p-Raf, p-PDK1, p-p90RSK, p-MKK3/6, p-MAPKAPK2) and overexpression level (OE), measured as GFP abundance. Because only phosphorylated species are observed, we added baseline offsets [p-ERK]_min_ and [p-MEK]_min_ , the minima along each reconstructed trajectory, to absorb unobserved baseline variation. To cover the full gradient we discretised overexpression into 50 bins, giving a dense family of trajectories from 0% to 100%. Each discrete time course was fitted with difference-of-logistic and exponential rise-and-fall forms under a smoothness regulariser coupling neighbouring bins, and differentiated analytically to give the target rates (Methods). We train on the fitted curves rather than the raw data, avoiding the noisy rate estimates that finite differencing would produce: they supply the rate target, the input trajectories and the initial condition, on a one-minute grid. Only one of the six measurements falls after 30 min, so we kept every grid point up to 30 min and a random 15 of the 30 later ones—46 per bin, identically for every model compared here; each of the three seeds redraws this sample as well as reseeding the search. The raw measurements are used only to score the result, at the six timepoints actually acquired. For physical consistency we added regularisation to the SR loss: penalties on excluding p-ERK (the state variable) or OE (the overexpression level), and on forms showing unbounded growth in p-ERK—locally linear terms with negative coefficients, or inversely linear terms with positive coefficients—so the recovered expression behaves as a self-consistent dynamics model under forward integration, not merely a fit to the ERK phosphorylation rate. An ablation showed that removing both stability penalties degrades predictive performance, although either penalty alone was sufficient (Fig. S1).

**Figure 4.**
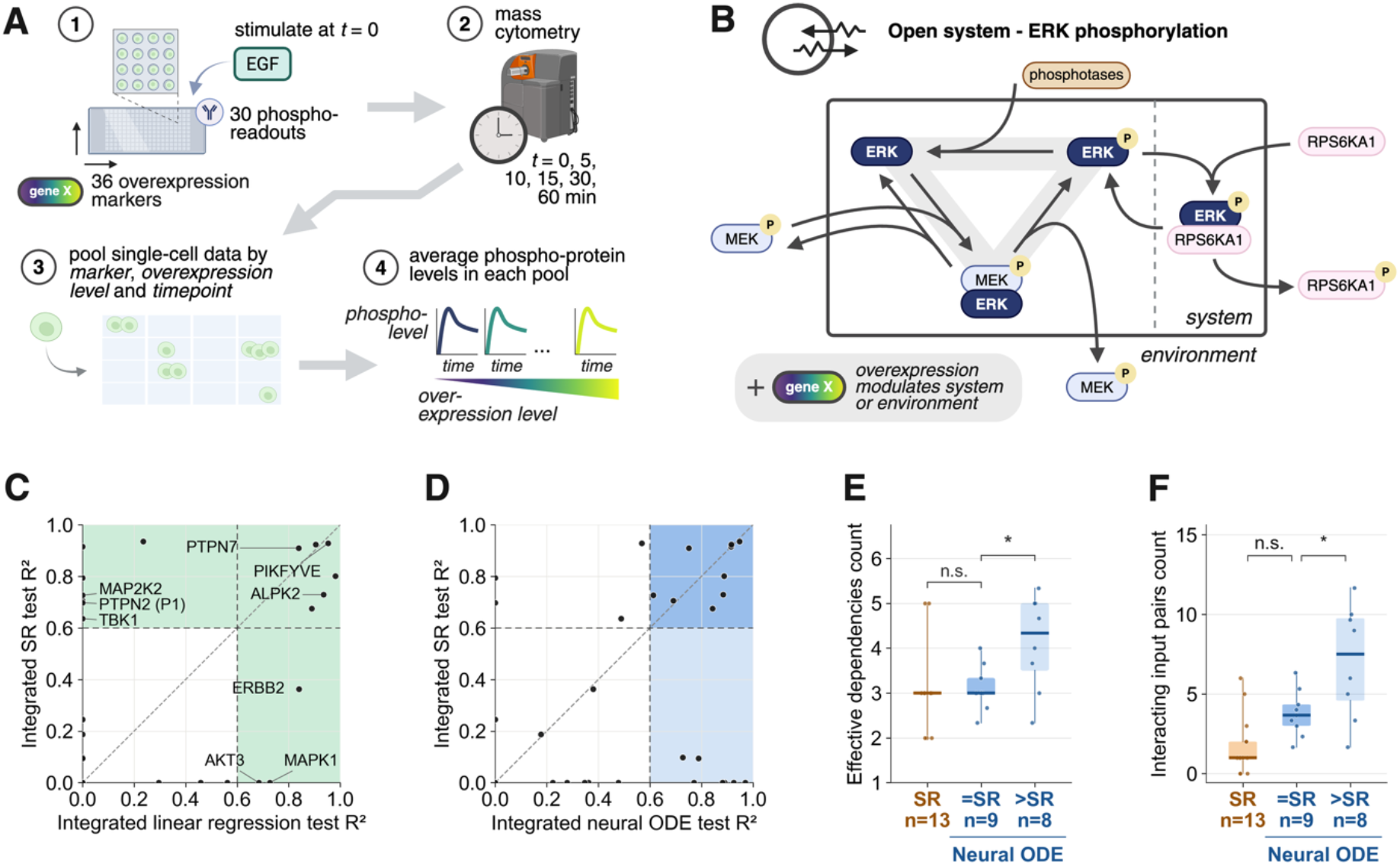
Workflow and evaluation of symbolic regression models for ERK phosphorylation across overexpression contexts. (**A**) Construction of overexpression-resolved trajectories from Lun *et al.*^51^. Cells were stratified by perturbation identity and GFP-based overexpression level, stimulated with EGF, sampled by mass cytometry at six timepoints, and pooled by marker, overexpression level and time; each trajectory is therefore a within-bin population average over the cells falling in one of 50 equal-count GFP quantile bins, not a tracked cell. (**B**) ERK-centred local signalling system. The black box marks the chosen system boundary; nodes crossing it are external inputs, outputs and perturbations. (**C**) Each point is one of the 40 contexts, by integrated linear-regression fit on all ten inputs (x) against integrated SR fit (y), the symbolic-regression fit using the seed selected by training R², both on the same test split; points above the diagonal favour SR. Contexts on the y-axis at x = 0 are those the linear model fails to extrapolate at all. All 40 contexts are plotted; only overexpression perturbations are labelled, the eight controls being shown unlabelled for legibility. (**D**) As (C), against integrated neural ODE fit, for the same 40 contexts. Threshold lines at R² = 0.6 mark satisfactory performance. The neural-ODE seed is chosen on a validation set and its test score reported, the SR seed by training R²; Neural ODE dependency counts are averaged over all three seeds. Only the Neural-ODE-success half is partitioned, since the panel asks whether the neural ODE also fails where SR fails: dark blue, solved by both (n = 9); light blue, neural ODE only (n = 8). (**E**) Effective dependencies where each method succeeds — the number of the ten model inputs whose mean absolute Jacobian exceeds 0.15 of the largest — for SR (n = 13) and for the neural ODE, the latter split by SR success (n = 9) or failure (n = 8). Neural-ODE counts are averaged over the three seeds; the SR count is that of the single retained law. Tick labels give each box’s comparison: =SR, contexts SR also solves; >SR, contexts the neural ODE solves and SR does not. (**F**) Interacting input pairs for the same three groups. In (E) and (F) boxes show the median and interquartile range, whiskers extend to the most extreme values not classified as outliers, and points are individual markers. Brackets: the neural ODE split by SR success (accuracy-residualised permutation test; *, p=0.025 in (E) and p=0.023 in (F); n.s., not significant).

We next asked whether these rate laws define a self-consistent dynamical model. We integrated each discovered ODE for p-ERK, treating the remaining signalling variables as time-dependent exogenous inputs, to reconstruct p-ERK time courses per context and overexpression level. All models were evaluated under an out-of-distribution split holding out the highest-GFP bins, so scoring reflects extrapolation to stronger perturbations than any seen in training. Every R² reported here, on test and training bins alike, compares the integrated trajectory with the raw measurements (Methods). PySR hyperparameters came from a 72-configuration grid search on a six-marker development set, selected by highest median training R² and applied unchanged to all 40 contexts (Methods and Fig. S3B). We used an integrated-trajectory R² of 0.6 as the criterion for satisfactory performance, since trajectories down to this cutoff still reproduced the main qualitative R² of 0.6 as the criterion for satisfactory performance, since trajectories down to this cutoff still reproduced the main qualitative features of the measured dynamics (Fig. S2). By this criterion SR met the threshold in 13 of 40 contexts (Fig. 4C; per-context results in Table S8), including a median of four of the ten model inputs. We compared this with a linear regression on all ten inputs (Fig. 4C, x-axis), evaluated on the identical split. SR reached the threshold on 13 contexts and the linear baseline on 9, with 6 solved by both (16 fit by either). In several of SR’s successes—including MAP2K2, PTPN2 (P1) and TBK1—the linear model does not extrapolate at all (R² = 0), so a linear combination of the same ten readouts carries no predictive dose response there and the gain is attributable to the non-linearity of the recovered law rather than to a better fit of the same form. Where the baseline wins instead—AKT3, ERBB2 and MAPK1—the symbolic fit ranges from partial to absent (R² 0.00–0.36). Across the 16 contexts either method fits (Fig. 4C, green), SR leads on 9 at a higher mean R² (0.67 vs 0.50). The two are also not matched in complexity: the baseline consumes all ten inputs, whereas the SR laws use a median of 4. This indicates SR is the more accurate coarse-grained model, and these markers require non-linear rate laws. Despite the density of the network surrounding ERK, non-linear coarse-grained models therefore exist and can be inferred directly from experimental data.

To test whether SR’s failures reflect the method or a genuine absence of a compact reduced description, we trained a neural ODE (NODE)^31^ with a group-sparsity penalty on its input Jacobian^53^ (seed selected on validation, penalty weight by calibration sweep; Methods, Fig. S3A, Table S14). Although far more expressive than SR, it is still forced to use as few inputs as it can without degrading the fit. To characterise structure across the large number of generated models, we used their Jacobians to identify non-negligible variable dependencies and their Hessians to identify non-negligible pairwise interactions among those variables, thereby characterising nonlinearity (Methods). Both quantities were averaged over three seeds and evaluated only for models that extrapolated successfully, since the sparsity structure of a poor fit is not interpretable.

Across the 40 contexts the NODE reached test R² > 0.6 in 17 and SR in 13, with 9 in common (Fig. 4D). Where SR failed, the NODEs relied on about one more measured variable (Fig. 4E; 4.12 vs 3.11; permutation p=0.025, adjusted for the NODE’s test accuracy) and carried about 86% more interacting pairs (Fig. 4F; 7.04 vs 3.78). One variable costs this much because nothing is separable: *k* inputs admit at most *k*(*k* − 1)/2 interacting pairs, and the measured pairs reach 96% of that maximum. A variable added to k others therefore couples to essentially all of them, adding k interaction terms rather than one. The pattern holds across all 40 contexts (4.09 vs 3.36; accuracy-adjusted permutation p=0.04), and the fraction of seeds recovering a generalising model falls as the count rises (Spearman ρ = −0.47, p=0.002). SR failure therefore largely reflects the absence of compact rate law rather than the failure to identify them.

Where a compact law is supported, we can ask what its nonlinearities mean. The variables and functional relationships SR selects—linking p-ERK dynamics to overexpression level and a sparse set of phospho-readouts—provide interpretable hypotheses about ERK self-regulation, how overexpression reshapes phosphorylation dynamics, and which pathway signals modulate the response. We illustrate with PIKFYVE and PTPN7, the two non-control markers with highest test R² for SR (0.93 and 0.91; PIKFYVE seed 44, PTPN7 seed 43). For PIKFYVE (Fig. 5A), OE enters positively, so increasing PIKFYVE raises the phosphorylation rate, while dividing the p-MEK drive by [p-ERK] makes ERK limit its own rate hyperbolically, with p-MAPKAPK2 entering as a separate negative term:

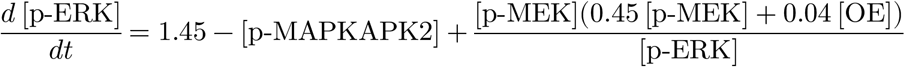

**Figure 5.**
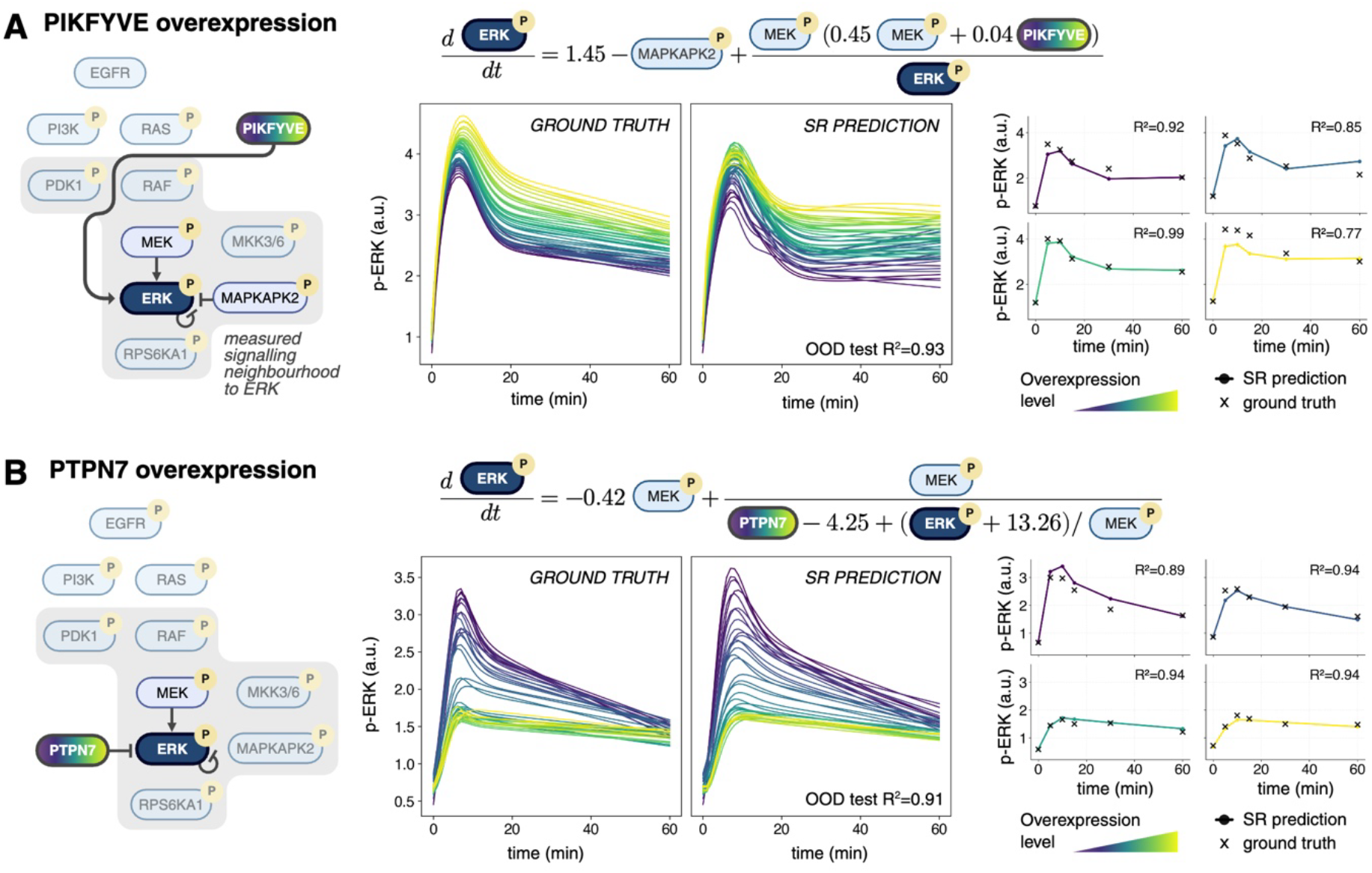
Representative overexpression contexts with highly predictive symbolic-regression models of ERK phosphorylation. (**A**, **B**) The two best-performing non-control contexts, each at the seed retained by training R²: PIKFYVE (seed 44) and PTPN7 (seed 43). Coefficients are shown to two decimal places; exact values are in Table S8. Left, the measured signalling neighbourhood around ERK, with the overexpressed marker and the variables retained by SR highlighted; the inferred rate law for *d*[*p* − *ERK*]/*dt* above. Centre, the fitted p-ERK trajectories — the parametric curves, not the raw readings —across the GFP bins retained by the fit-quality filter (49 for PIKFYVE and 39 for PTPN7, of 50; Methods), beside trajectories from integrating the inferred model; curve colour encodes overexpression level. Right, four representative overexpression-level groups spanning the gradient, with crosses for the raw p-ERK measurements, lines for integrated predictions, and the per-trajectory R² of each bin shown; the three lower panels are training bins and only the highest-GFP panel is held out.

Receptors continue to signal to ERK after internalisation^54^, and PIKFYVE controls how endosomes mature^55^, so it plausibly sets the strength of the p-MEK drive rather than acting on ERK itself. Here p-MAPKAPK2 serves as a readout of p38 activity, and p38 induces DUSP-family phosphatases that dephosphorylate the ERK activation loop, so a negative coefficient is the expected sign^56^; p38 signalling has separately been linked to PIKFYVE function^57^. For PTPN7 (Fig. 5B) the law needs no exogenous driver beyond p-MEK:

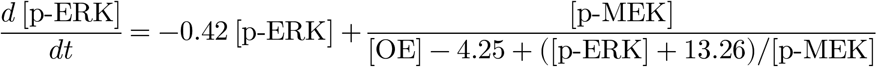

a saturating p-MEK drive against p-ERK-dependent removal. Overexpression enters only in the denominator, through [OE] − 4.25, so increasing PTPN7 monotonically suppresses the rate. This agrees with known biology: PTPN7 binds ERK through a kinase-interaction motif and dephosphorylates its activation loop, and its overexpression blocks MAPK activation^58^. These results show that SR both reconstructs time-resolved trajectories with high fidelity and identifies how specific overexpressions rewire signalling logic, establishing it as a route from high-dimensional measurements to compact dynamical laws that are predictive and mechanistically legible—and, where no such law is supported, as a data-driven probe that suggests the dynamics are not compressible.

## Discussion

Coarse-graining is unavoidable in modelling intracellular signalling^1^. We show that, done in a data-driven way, it can be used not simply as an implicit step in model construction, but as a tool for inferring scale- and context-relevant dynamical laws. Across increasingly realistic systems, SR proved both a method for coarse-grained model discovery and a data-driven probe of coarse-grainability: it tests whether a self-contained reduced model is supported by the measured variables, and recovers structured, mechanistically informative equations when it is.

We first showed that SR can recover the Michaelis-Menten equation from synthetic data generated under the idealised conditions in which it is valid. PySR outperformed the other methods tested, likely because it handled targets spanning multiple orders of magnitude more naturally. Although rediscovering known laws is a standard SR benchmark^37^, enzyme kinetics is a useful test case within a broader class of nonlinear saturating relations—Michaelis-Menten and Hill-type laws—that recur across biological domains.

PySR also recovered Michaelis-Menten after relaxing those assumptions through product inhibition and system openness, indicating that SR can recover literature laws beyond their strict derivation regime. This suggests that Michaelis-Menten behaves as a robust emergent description of enzymatic activity rather than merely a narrow special-case approximation, consistent with recent work on the universality of Hill-type input–output laws^59^. By contrast, enzyme dimerisation was a genuine mechanistic departure, introducing association-dependent activation and unmeasured species; there the observed variables no longer appeared to support a low-complexity, self-contained law at the chosen scale—a failure traceable to a specific missing information rather than to the search.

Applied to experimental phospho-proteomic data, we asked whether the coarse-grained models of ERK phosphorylation were accessible from the available measurements. In a subset of overexpression contexts they were, and the recovered laws were interpretable and biologically meaningful. Elsewhere SR failed, and across those contexts the sparse neural ODE needed on average about one additional input and nearly twice as many interacting pairs than where SR succeeded. Together, these results suggest a common limitation: as the measured variables become less adequate for closing the dynamics, a flexible model must represent them through increasingly dense coupling, until even the neural ODE fails. This is supported by the observations elsewhere that either transformation^60^, or augmentation with latent states ^5,6^ are necessary to find sparse rate laws. Indeed, Mori-Zwanzig theory^61,62^ formalises this possibility, showing that eliminating unobserved species generically introduces memory into the reduced dynamics, which neither model can represent. Such failures therefore suggest that no coarse-grained description exists in the observed variables, motivating a different set of measurements, coordinates or scale—or models incorporating latent states.

Where successful, SR identified predictive, mechanistically interpretable ODEs yielding concrete hypotheses for how overexpression rewires ERK signalling. The two exemplars mark the range of what this yields: for PTPN7 the recovered law recapitulates an activity already established for the enzyme, so it serves as a check that the method recovers real biology rather than as a discovery; for PIKFYVE it instead points to an effect on the p-MEK drive rather than on ERK itself, for which we are aware of no direct evidence, and which is therefore a testable prediction of the kind this approach is meant to generate. Across these cases, SR’s recovered laws were consistent with the imposed stability constraint on p-ERK and explained condition-specific shifts in ERK behaviour with parsimonious models using only a small subset of phospho-protein readouts.

Because phospho-signalling markers in local network neighbourhoods are strongly correlated^51^, alternative fits from different seeds sometimes exchanged which phospho-inputs appeared explicitly. This reflects a limitation of the observations rather than necessarily of the method: multiple correlated measurements can support similarly predictive reduced models of the same behaviour. The inferred interactions should therefore be interpreted as effective dynamical dependencies rather than direct molecular interactions. Additional perturbational data may resolve which correlated variables are most directly involved. Even so, because the models are dynamical, an included term implies directed predictive influence on future p-ERK behaviour beyond p-ERK’s own history, in a Granger causality-like sense^63^. They therefore achieve the goal of mechanistic modelling here: compact, interpretable hypotheses for how specific overexpressions reshape ERK phosphorylation dynamics.

A limitation of the study lies in the data rather than the method. The overexpression gradient reaches extreme levels at its upper end, far beyond physiological abundance, where induced interactions may not reflect signalling at native levels—so dependencies recovered from the high-expression bins may be specific to that regime. Combined with a large underlying network and a highly diverse intervention set^51^, this makes a single unified coarse-grained model, well constrained across all conditions, difficult to infer. We therefore infer perturbation-specific interactions rather than enforcing a shared global structure.

We also focused on one state variable, p-ERK, treating other phosphoproteins as observed exogenous inputs. Extending this to multi-state dynamics is considerably harder: one must infer a coupled system whose joint integration is identifiable, numerically stable and dynamically coherent. Current SR tools such as PySR are not designed for this: they infer one static equation at a time and cannot enforce consistency across states^36^. Methods for multi-equation dynamics discovery such as SINDy^40^ offer a complementary direction but require a predefined function library. We did not evaluate SINDy here: our synthetic closed-system benchmarks did not provide derivatives in the required form, and the diversity of ERK phosphorylation laws PySR recovered would be hard to capture in a fixed library. A key future direction is therefore to combine free-form SR’s expressivity with system-level dynamical constraints for learning coupled ODE systems. SR could also identify recurring motifs to inform targeted SINDy libraries. Universal Differential Equations^34^ are another avenue, with SR providing a compact mechanistic backbone and neural terms capturing residual context-specific dynamics.

More broadly, rather than prespecifying the effective laws expected to govern signalling dynamics, future studies could use SR to ask which low-dimensional laws a perturbational dataset supports, where they fail, and which components of signalling state are missing when they do. This creates a concrete opportunity for experimental design: where compact laws hold, the inferred terms generate testable hypotheses about the dependencies and nonlinearities that implement rewiring; where they fail, the mismatch points to hidden state and motivates targeted additions to the measurement panel or perturbation scheme. SR could thus accelerate the model-experiment loop at the heart of mechanistic biology, helping identify which features of a complex signalling system are essential for explaining its dynamics.

## Materials and Methods

### Synthetic enzyme systems

Mass-action ODE models of two-, three- and four-step catalysis, closed and open. The three-step adds reversible enzyme–product binding; the four-step adds dimerisation, making activity depend on an association state not fully exposed in the observables. Rate constants and initial conditions sampled log-uniformly to span kinetic regimes, trajectories integrated to steady state, observables recorded under the standard or total quasi-steady-state lens (SI Appendix).

### Data degradation

Five degradations of the open three-step dataset: measurement noise, training-set size, departure from Michaelis–Menten kinetics, restriction to defined kinetic limits, and choice of observable. The first three were each crossed with the free- and total-substrate lenses. Levels in SI Appendix.

### Experimental data

Published time-resolved phospho-proteomic measurements of ERK signalling across 40 overexpression contexts. Acquisition is by mass cytometry snapshots, so trajectories are within-bin population averages: cells binned into 50 GFP quantiles, p-ERK fitted per bin with a parametric rise-and-fall model under a neighbour-coupling smoothness penalty. The fit supplies the rate target (its analytic derivative), the initial condition and the exogenous inputs; scoring is against the raw measurements. Inputs were p-ERK, p-MEK, p-Raf, p-PDK1, p-p90RSK, p-MKK3/6, p-MAPKAPK2, two baseline offsets and GFP.

### Out-of-distribution split

The highest 20% of GFP bins withheld per context, so models are scored on extrapolation beyond trained overexpression levels. The neural ODE also withheld the next-highest 20%, immediately below the test block, for early stopping. Both partitions are deterministic; seeds vary only search initialisation.

### Symbolic regression

PySR with the four arithmetic binaries, so expressions stay interpretable as rate laws; total-substrate analyses also allowed squaring and square root. Synthetic fits used a log-space loss, targets spanning many orders of magnitude. Experimental fits added soft penalties on omitting p-ERK or GFP and on divergence under integration (ablated, Fig. S1). Search settings and the 72-configuration sweep in SI Appendix (Table S9).

### Baselines

Synthetic: the analytical Michaelis–Menten law and a multilayer perceptron, with AI-Feynman, DSO and KAN benchmarking alternative SR implementations. Experimental: linear regression on all ten inputs—not complexity-matched, by design—and a sparse neural ODE. Architectures and the 54-configuration sweep in SI Appendix (Table S10).

### Metrics

Synthetic accuracy is the median relative absolute error. Experimental performance is the R² of the ODE-integrated p-ERK trajectory against the raw measurements at the six acquired timepoints, taken as the median across test bins and floored at zero, so a reported 0.000 denotes R² ≤ 0. For symbolic regression the seed with the highest training R² was retained per context and its test score reported; for the neural ODE the seed with the highest validation R² was retained and its test score reported, so neither selection uses test data. All statistical tests are two-sided and treat the context as the unit of analysis; reported p-values are unadjusted and these comparisons are exploratory and hypothesis-generating. Effective dependencies are the number of inputs whose mean absolute Jacobian exceeds 0.15 of the largest, applied to the neural ODE averaged over three seeds, and the recovered model on its best seed; interacting pairs are off-diagonal input-Hessian entries above 0.25 of the largest among the selected inputs. Neither cut is critical: the contrast holds for Jacobian thresholds from 0.05 to 0.25 (Table S12) and for off-diagonal thresholds from 0.05 to 0.30 (Table S13), and we report where it fails. Both models’ seeds are selected on data disjoint from the test split, while quantities such as complexity are averaged over the three seeds. Complexity contrasts are exact permutation tests on the difference in group means, computed on counts residualised on the neural ODE’s held-out R² so a difference in measurement quality cannot present as a difference in complexity.

### Reproducibility

Analyses used fixed seed 42 unless stated; degradation and experimental runs used seeds 42–44 (SI Appendix).

## Data availability

The mass cytometry dataset underlying Lun *et al.*^51^ is publicly available via Mendeley Data (DOI: 10.17632/3kh7ypz232.1). Synthetic datasets generated from mechanistic models can be reproduced using the provided Snakemake workflow and generation scripts.

## Code availability

All code to reproduce the synthetic simulations, SR training, degradation experiments and experimental ERK analyses is available at [n/a] and executable via Snakemake. Full hyperparameter grids and per-context metrics are deposited there.

## Acknowledgements

We thank the whole Dynamics of Living Systems lab for helpful discussions and feedback on the manuscript.

## Funding

F.F. and T.P. were supported by the Francis Crick Institute, which receives its core funding from Cancer Research UK (CC2242), the UK Medical Research Council (CC2242), and the Wellcome Trust (CC2242), as well as the European Union (ERC, DeepMechanism, grant no 101163005).

## Competing Interests

F.F. reports consulting honoraria from Deep Origin, but this had not influence on the study.

## SI Methods

### Synthetic enzyme systems

#### Two-step mechanism

We considered enzyme-catalysis models in which a substrate *S* is converted into a product *P* by an enzyme *E* through formation of an enzyme–substrate complex *ES*.

#### Closed system

In the closed setting, species evolve only through the reaction itself, with no exchange with the environment. Our goal was to recover the coarse-grained catalytic law rather than to model the full product trajectory explicitly. We therefore took the instantaneous catalytic rate, *v*(*t*) = *dP* /*dt* = *k*_cat_[*ES*], as the quantity of interest. We further assumed rapid equilibration of enzyme-substrate binding relative to catalytic turnover, so that [*ES*] is effectively determined by the current enzyme and substrate abundances. Because the product does not feedback on the dynamics in this minimal closed system, explicitly introducing *P* as a dynamical state is unnecessary for the rate-law recovery task. Accordingly, we omitted the product state from the simulated system and represented catalysis as depletion of the enzyme-substrate complex with release of free enzyme and substrate, while taking *v*(*t*) = *k*_cat_[*ES*] as the coarse-grained readout. The reaction scheme is therefore

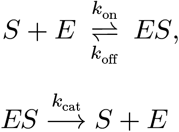

with governing equations derived from mass-action kinetics:

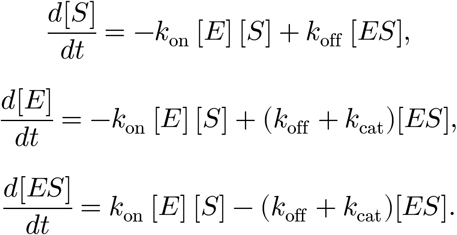

#### Open system

In the open setting, we introduced explicit product formation, reverse conversion, and enzyme turnover. The reaction scheme is

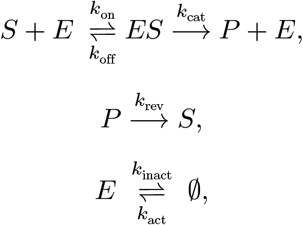

with governing equations

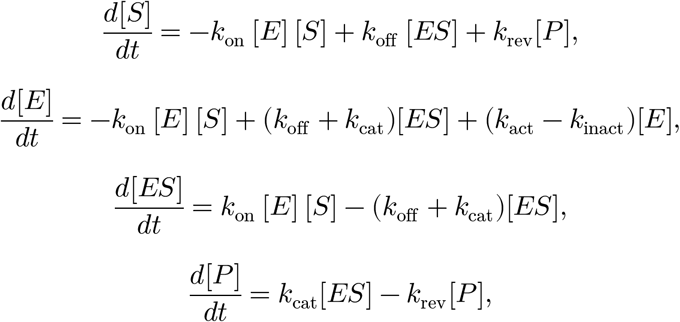

where the nominal enzyme level is set by the balance of activation and inactivation, *k*_act_ = *K*_0_*k*_inact_, so that in the absence of substrate the enzyme relaxes to the baseline level *E* = *K*_0_.

#### Three-step extension

This mechanism extended the two-step scheme by allowing reversible product binding to the enzyme,

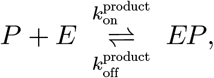

thereby introducing an inhibitory product-bound complex *EP* and relaxing the assumption that catalysis proceeds independently of accumulated product. In the open system, where we explicitly model the product during catalysis, the catalytic step becomes

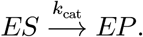

#### Four-step extension

This mechanism further introduced reversible enzyme dimerisation,

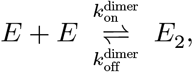

alongside substrate binding, catalysis, and product inhibition, with the dimer *E*_2_ acting as the catalytically active enzyme species. Activity therefore depended on association-dependent enzyme state rather than enzyme abundance alone. In this setting, dimer species were present in the mechanistic simulation but not fully exposed in the reduced observables, creating a restricted-observability regime in which the measured variables need not uniquely determine the effective rate.

### Synthetic data generation

#### Parameter sampling and dataset generation

For each train, validation and test split, kinetic parameters were sampled independently in log_10_-space as

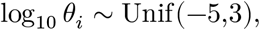

for each sampled parameter θ*_i_*, before conversion to natural-log scale for simulation. We used 2^16^, 2^14^ and 2^14^ samples for the train, validation and test sets, respectively. In the dynamic setting, additional resampling constrained enzyme inactivation relative to reverse conversion to avoid extreme timescale separation and sampled the nominal enzyme pool *E*_0_ within a bounded factor around its pre-equilibrium scale.

#### Simulation schedule and steady-state criterion

Closed-system models used a terminal steady-state readout only, whereas open-system datasets used a two-phase AMICI^64^ simulation. First, each condition was first pre-equilibrated to steady state. Second, the enzyme abundance *K*_0_ was resampled in, with *K*_0,pre_/*K*_0,post_ sampled log-uniformly as

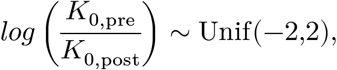

while all other fixed parameters (including *k*_inact_ and *k*_rev_) were unchanged between phases. The system was then simulated again until equilibrium and sampled at *t* = 0, a 20 timepoint log-spaced grid over [*t_min_*, *t_max_*], and a terminal steady-state readout. The interval [*t_min_*, *t_max_*] was calibrated to ensure that the simulated dynamics both departed from and returned close to equilibrium over the observed window. Steady state was defined using AMICI’s internal Newton-step criterion: for state vector x, vector field *f*(*x*), Jacobian *J*(*x*), and Newton correction Δ*x* satisfying *J*(*x*)Δ*x* = −*f*(*x*), steady state is accepted when,

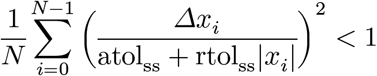

using atol_ss_ = 0 and rtol_ss_ = 10^−8^, and where is *N* is the number of states.

#### Quality control and resampling

Simulations were discarded and resampled whenever numerical integration failed or any required state or derived quantity was non-finite. Formally, a simulation was retained only if

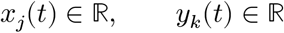

for all required states *x*_j_, derived observables *y_k_*, and sampled timepoints *t*, and likewise after any downstream log transformation where applicable.

#### Preprocessing and splitting for symbolic regression

Raw train/validation/test datasets are produced directly during generation. A subsequent preprocessing step merges raw splits and constructs processed datasets, performing an 80/20 split into train/test. In the dynamic setting, SR operates on pooled trajectories and is not trajectory aware.

### Experimental data

#### Dataset and preprocessing

We used time-resolved perturbational phospho-proteomic measurements from Lun *et al.*^51^. The dataset comprises 40 perturbation contexts: 32 overexpressed constructs and 8 controls. Trajectories are indexed by marker, GFP perturbation intensity, and timepoint. Preprocessing discretises perturbation intensity into 50 bins per marker using quantile binning. Replicates are averaged by grouping on (marker, GFP bin, timepoint). Bins with less than 5 measured timepoints are excluded from parametric fitting.

#### Parametric trajectory fitting and derivative estimation

To obtain smooth rate targets, we fit per-overexpression marker phosphoprotein trajectories using two candidate parametric forms:

1. Rise-and-fall model: 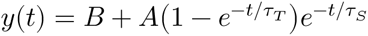
2. Difference-of-logistics with tail: 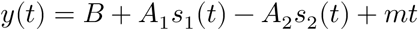 with logistic terms 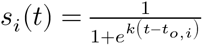

Fits are performed jointly across all eligible GFP bins for a given marker by minimising a robust least-squares objective (trust-region reflective, Huber loss, max_nfev=1000) with a neighbour-coupling smoothness penalty on bin-wise parameter vectors, implemented as residual terms 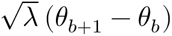 for adjacent bins (with *λ* set to 1.0). Model selection prefers the highest global coefficient of determination

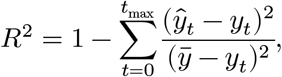

breaking near-ties within 1×10^-3^ by lowest Akaike information criterion with a correction for small small sizes (AICc). Analytic derivatives of the selected fit are computed and stored as derivative targets. Bins with p-ERK fit quality below a threshold R^2^=0.95 are excluded to ensure the derivatives are representative of the true underlying dynamics. To densify trajectories, the pipeline generates per-minute dense trajectories by evaluating fits on a 1-minute grid. GFP values at added timepoints are filled using per-bin mean GFP.

#### Train/test splitting for ERK models

Experimental models are evaluated on splits disjoint in GFP bins. With bins sorted by overexpression, the highest 20% form the test set and the remaining 80% the training set; the neural ODE takes a third block, the next-highest 20%, for early stopping, leaving 60% for training (10, 10 and 30 of the 50 bins per context). Both cuts are deterministic. The seed varies three things: model initialisation, the symbolic search, and a subsample of the dense grid. Only one of the six measured timepoints falls after 30 min, so the rest of that window is interpolation; every grid point up to 30 min is therefore retained and 15 of the 30 later points are drawn at random per marker and GFP bin, giving 46 of the 61 points per bin. Every model fitted to these data uses the same draw, so all are trained on identical points. Evaluation therefore reflects out-of-distribution extrapolation across overexpression intensity.

### Symbolic Regression

#### Task for synthetic settings

For the synthetic enzyme-system data, we used the SR methods to infer a coarse-grained expression for the enzyme-substrate complex abundance,

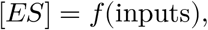

across the two-step, three-step, and four-step enzyme mechanisms under both closed- and open-system conditions. The default input set comprised the reduced variables and kinetic parameters exposed at the coarse-grained level. For the two-step benchmark, these inputs were *S*, *E*_tot_, *k*_cat_, *k*_off_, and *K_D_*. For the open system setting, the input set was expanded to include *k*_inact_, while remaining restricted to same observables available at the reduced level. The regression target was the enzyme-substrate complex abundance [*ES*], chosen as a common catalytic observable across all synthetic systems. In the classical two-step mechanism, the catalytic flux rate is given by

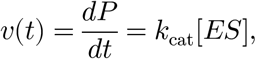

and, under the quasi-steady-state approximation, this theoretically yields the Michaelis– Menten rate law. In the extended three-step and four-step mechanisms, [*ES*] was retained as a consistent coarse-grained observable, although it does not in general define an exact reduced flux law on its own, due to added mass-action terms on the right-hand side of the equation above.

#### Task for experimental settings

SR was applied to infer a coarse-grained rate law for p-ERK:

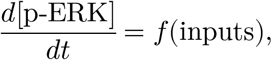

where the default input set includes GFP, p-ERK1-2, and local signalling variables: p-MEK1-2, p-Raf, p-p90RSK, p-MAPKAPK2, p-PDK1, and p-MKK3-6, together with two baseline offsets (p-ERKmin and p-MEKmin), giving ten inputs in total. The SR target was the estimated time derivative of p-ERK using the analytical derivative of the relevant fitted parametric model.

#### PySR configuration

PySR was used for all synthetic analyses beyond the initial benchmark and for all experimental ERK analyses. In the standard synthetic setting, PySR used the binary operator set {+, −,×,÷}, with no unary operators (except the tQSSA analyses; see below). Search was performed with: niterations=350, population_size=30, populations=15, maxsize=20, parsimony=1. To account for the large dynamic range of the synthetic targets, PySR was trained with a custom log-space elementwise loss,

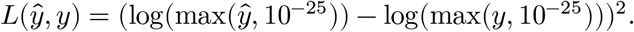

This loss penalises multiplicative error rather than absolute deviation and was important for accurate recovery of rate laws across regimes spanning many orders of magnitude. Runs used batching and annealing and were executed deterministically in serial mode. For each PySR run, candidate expressions were selected using the default PySR accuracy-complexity trade-off score.

For the tQSSA measurement-lens analyses, PySR used the same general configuration, except that: the unary operator library was extended to include the square root and squaring operators and the maximum expression size was increased to maxsize=45. These changes were introduced because the total-substrate lens induces more algebraically complex reduced forms than the standard Michaelis–Menten setting.

For the experimental ERK analysis, PySR used the same general search framework but with the default PySR linear-space element-wise mean squared error loss augmented with custom stability penalties (see below). The following search parameters were increased: niterations=1400, populations=30, population_size=30, maxsize=26, parsimony=0.8 (selected by the sweep described under Hyperparameter tuning).

#### Configuration of alternative SR methods

The non-PySR SR methods were used only in the initial synthetic benchmark. These methods do not natively support the kind of elementwise custom loss that PySR has, so they were trained using their standard objectives with the log-transformed target, while still using linear-scale inputs (except KAN, whose inputs were kept on the log scale, where its spline basis is better conditioned). Logarithmic operators were included in their expression libraries when possible so that log-like structure could still appear in learned formulas. All benchmarked methods were run under a fixed training-time budget of 10,800 seconds (3 h) to reflect practical usability in the broader workflow.

AI-Feynman was run with the operator string “+-*DL”, corresponding to addition, subtraction, multiplication, division, and log. Internal search settings followed wrapper defaults.

DSO used the function library specified by the pipeline configuration; in the log-output setting this effectively corresponded to {add,sub,mul,div,log}. The model used a least-squares regression head with Adam optimisation and entropy regularisation, with total training samples determined by the configured batch size and iteration count.

For the KAN-based symbolic baseline, we first fit a spline-based KAN and then converted the trained network into an explicit symbolic expression using an automated symbolification step. Each learned one-dimensional spline function was approximated by the best-matching analytic form from the library {*x*, *x*^2^, *x*^3^, 1/*x*, log}, thereby biasing the extracted expressions toward low-order polynomial, reciprocal, and logarithmic dependencies. This baseline differs conceptually from tree-search SR methods in that it learns a flexible neural representation first and only subsequently extracts a symbolic approximation.

Full implementation details for these benchmark baselines are provided in the repository configuration.

#### Sample sizes used in the SR synthetic benchmark

Following method-specific best practices, we used different effective training sample counts for the initial four-method benchmark: PySR: 3000 samples, AI-Feynman: 5000 samples, DSO: 38400 samples, KAN: 4000 samples. These differences reflect method-specific computational requirements rather than strict equivalence of optimisation budget.

#### PySR penalty loss terms for experimental settings

For the ERK-signalling task, predictive accuracy alone was insufficient: candidate rate laws also had to remain interpretable and numerically stable when integrated as one-dimensional ODEs. We therefore used a custom PySR loss written in Julia that augmented the data-fitting objective with structural penalties. These included: (i) enforcing dependence on p-ERK and GFP, (ii) discouraging near-zero sensitivity to these variables, and (iii) penalising locally unstable functional forms with respect to p-ERK by detecting approximately linear dependence in p-ERK or 1/p-ERK directions via finite-difference sensitivity tests and applying wrong-sign penalties (penalty magnitudes and tolerances defined in the script; default penalty scale 1000 and dependence tolerance 10^-4^).

### Baseline methods

#### Michaelis-Menten baseline (synthetic degradation experiments)

As an analytical baseline, we evaluated the standard Michaelis–Menten rate law on the same synthetic test sets used for SR in the degradation experiments.

#### Multi-Layer Perceptron baseline (synthetic degradation experiments)

A neural baseline MLP was implemented and trained on the same inputs and train/test splits as for the synthetic SR setting within each synthetic regime. Inputs were standardised on the training set. Hyperparameters were selected by a separate tuning procedure on validation performance, after which all results use a fixed architecture with two hidden layers (256 and 128 units) with SiLU activations and dropout (0.3). The model was trained for up to 600 epochs with batch size 256 using RMSprop (learning rate 6.53×10^-4^, weight decay 1.35×10^-4^), with gradient clipping (global norm 4.0) and a cosine annealing learning-rate schedule (CosineAnnealingLR). Model selection used a 10% validation split evaluated every 10 epochs, with early stopping (patience=70, min_delta=1×10^-4^); an independent 10% test split was held out for reporting, and all regimes used at least 10000 samples (except in dataset size reduction experiment).

#### Linear regression baseline (experimental)

As a reference model we fitted, per overexpression context, a linear regression of on all ten model inputs *d*[p − ERK]/*dt*, trained and evaluated on the same out-of-distribution split and scored by the same integrated-trajectory R^2^.

#### Sparse neural ODE baseline (experimental)

We trained a per-marker sparse neural ODE with a small MLP right-hand side (hidden_dim = 64, 4 hidden layers, tanh activation), integrated by an adaptive Dopri5 solver with linear interpolation of the exogenous inputs. p-ERK1-2 was the dynamical state; the remaining phospho-markers and GFP were time-dependent exogenous inputs. Training minimised a trajectory-level MSE (not derivative MSE) by backpropagation through the solver, using Adam (lr = 3×10^-3^, weight decay 10^-5^), batch size 64, up to 200 epochs with early stopping on validation loss (patience 20). To encourage reliance on a compact set of first-order dependencies we added a group-sparsity (L21) penalty on the columns of the input Jacobian evaluated at the training inputs, weighted by λ = 3.0. The seed with the highest validation R² is retained and its test R² reported.

### Metrics

#### Symbolic complexity

Symbolic complexity was computed using the default PySR complexity metric, in which the complexity of an expression is defined recursively over its expression tree as the sum of the complexities of all constituent nodes, with each operator, constant, and variable occurrence assigned unit complexity. Thus, for an expression tree *T*,

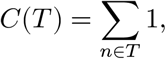

where the sum runs over all nodes n in the tree, including operators, constants, and variable leaves.

#### Synthetic evaluation

Synthetic evaluation uses the median relative absolute error (MdRAE) metric where

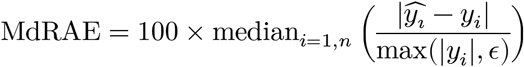

#### Experimental evaluation

Performance is summarised by binwise R^2^ statistics computed at the experimentally acquired time points, across GFP bins (median). The comparison is against the raw binned measurements, not the parametric fit that supplies the model’s target and inputs; scoring against the fit would compare a model with a smoothed version of its own supervision. The same target is used for symbolic regression, the linear baseline and the neural ODE. Downstream plotting and summaries use an R^2^ threshold of 0.6 to denote high-performing models.

#### Effective dependency count

For each trained neural ODE we compute the mean absolute Jacobian across the ten inputs and count those exceeding 0.15 of the largest, averaging over three seeds; interacting pairs are off-diagonal input-Hessian entries above 0.25 of the largest among the selected inputs.

#### Synthetic data degradation experiments

All degradation experiments used the open-system three-step model as their testbed, repeated runs across three seeds and used datasets of 3000 samples for each degradation condition.

#### Measurement noise regimes

Noise experiments add Gaussian noise to the target column only in log space:

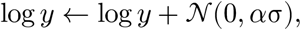

followed by exponentiation back to linear scale. σ is the dataset’s standard deviation and noise multipliers were *α* ∈ {10^−2^, 10^−1^, 1}, alongside a no-noise condition.

#### Deviation-from–Michaelis-Menten regimes

Deviation regimes quantify per-row deviation as the absolute log error between the mechanistic target and a Michaelis-Menten form predictor computed directly from features (no fitting). Rows are binned into thresholds: low: < 0.01, medium: 0.01–0.1, large: 0.1–1.0, very large: ≥ 1.0; and SR is trained/evaluated separately within each bin.

#### Kinetic-limit regimes

Kinetic regimes form subsets based on ratios computed from dataset columns, namely the substrate-to-enzyme proxy: *S*/*E_tot_* and the Michaelis ratio: *K_M_* /*S*, using *K_M_* = (*k*_off_ + *k*_cat_ + *k*_inact_)/(*K_D_ k*_off_). Thresholds are used to define extreme regimes (ratio ≥1000 or ≤0.001), and rows are filtered accordingly.

#### Alternative measurement lens (tQSSA shift)

To emulate measuring total substrate rather than free substrate, the tQSSA variant replaces the substrate feature with *S*_tot_ = *S* + *ES*, implemented as an in-place feature overwrite. An analytic tQSSA baseline predictor is used for comparisons where applicable.

#### Numerical integration

To assess whether inferred rate laws define a self-consistent dynamical model, we numerically integrated the discovered ODE for p-ERK, treating other signalling variables and GFP as time-dependent exogenous inputs. We perform ODE-solver integration using diffrax (Kvaerno5) with tolerances ODE_ATOL=1×10^-6,^ ODE_RTOL=1×10^-4,^ step constraints (ODE_DTMAX=1.0), and a large step budget (max_steps=200000). Exogenous inputs are interpolated in time in the integration grid, while p-ERK is supplied from the evolving state.

### Hyperparameter tuning

#### PySR (experimental)

PySR hyperparameters were chosen by a grid sweep of 72 configurations — max-iterations ∈ {700, 1400}, populations ∈ {30, 60}, population size ∈ {30, 60}, maximum expression complexity ∈ {18, 22, 26}, parsimony coefficient ∈ {0.1, 0.8, 3.0} — with binary operators fixed to {+, −, ×, ÷}. Each configuration was run on a six-context development set (AKT3, ALPK2, DYRK2, ERBB2, PIKFYVE, PTPN7) with three seeds (42, 43, 44), giving 1,296 fits. We selected the configuration with the highest median integrated R^2^ on the training GFP bins, so that the test highest-dose bins played no part in configuration choice: 1400 iterations, 30 populations, population size 30, maximum complexity 26, parsimony 0.8 (median training R^2^ 0.888, ranked first of 72; Table S9). This configuration was applied unchanged to all 40 overexpression contexts and used for all reported results. The arms were ranked against the parametric fit rather than the raw measurements; on the training bins the two agree to a median per-bin R² of 0.99.

#### Multi-Layer Perceptron baseline (synthetic degradation experiment)

We implemented an Optuna-based tuner to select MLP hyperparameters by minimising test test MAE (with targets in log space). Studies used Optuna’s TPE sampler (TPESampler) and ran for up to 100 trials (or an optional timeout). Each study constructs a single train/val/test, then reuses that split across trials while varying only network/training hyperparameters. The search space comprised: learning rate (log-uniform 1×10^-5^–1×10^-1),^ batch size {128, 256, 512, 1024}, hidden-layer templates {[32], [64,32], [128,64], [256,128], [512,256,128], [1024,512,256,128]} with associated dropout hints, dropout rate (0.0–0.5 step 0.05 when not overridden), weight decay (log-uniform 1×10^-7–^1×10^-1),^ activation {ReLU, SiLU, Tanh}, optimiser {Adam, AdamW, RMSprop}, scheduler {None, ReduceLROnPlateau, CosineAnnealingLR, ExponentialLR}, early-stopping patience (20–80 step 10), and gradient-clip norm (0.0–5.0 step 0.5).

#### Sparse neural ODE

Architecture and optimisation settings were selected on the model without sparsity regularisation: hidden dimension {32, 64, 128}, hidden layers {2, 3, 4}, learning rate {10^-3^, 3×10^-3^, 10^-2^}, activation {tanh, softplus}, evaluated on in-distribution validation R² with a single seed (Table S10). We chose the final configuration (hidden_dim = 64, 4 layers, learning rate 3×10^-3^, tanh) using in-distribution validation only.

#### Sparsity regulariser calibration

Using the selected configuration from the sweep we calibrated the sparsity regularisation term. We chose the group-sparsity (L21) penalty on the columns of the input Jacobian. Its strength was set by a one-dimensional sweep over the regularisation parameter in {1, 2, 3, 5, 8, 15, 30} using an elbow rule. The largest parameter value within 0.05 of the best mean in-distribution validation R2 was 3.0 (Fig. S3A).

---

**Figure S1.**
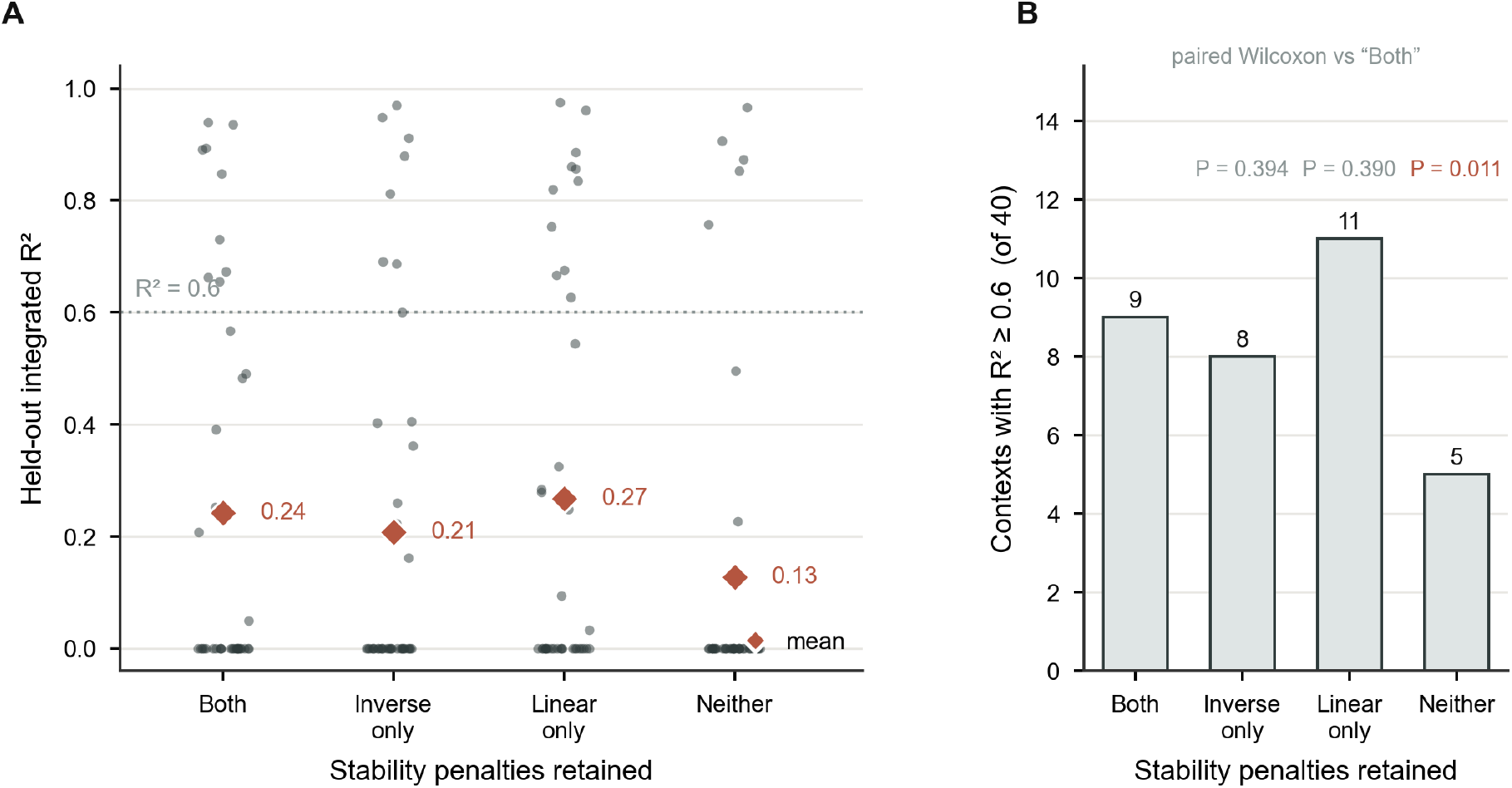
Ablation of the custom symbolic-regression loss penalties. Each of the four loss configurations was refitted on all 40 overexpression contexts (seed 42) under the selected configuration and the top-GFP-bin out-of-distribution split, so the values are on the same scale as every other experimental result reported here; the full-penalty arm reproduces the frozen run exactly (40/40 identical rate laws). (A) Test integrated R² per context. Most contexts score 0 under extrapolation, so the median is 0 in every configuration and the mean (diamond) is the informative summary. (B) Number of contexts clearing R² = 0.6, with paired Wilcoxon signed-rank tests against the full-penalty arm across the 40 shared contexts. Removing both penalties degrades performance (mean R² 0.24 → 0.13, 9 → 5 contexts above threshold, p=0.011), but removing either one alone does not (p=0.39 in both cases). Since the difference in performance between choosing one only or both penalties isn’t statistically significant, we opt for both out of precaution. Counts are for seed 42 alone, so the full-penalty total of 9 differs from the 13 of 40 reported in the main text, which selects a seed per context. Contexts whose inferred law fails to integrate score 0 rather than being dropped, so configurations are not graded only on the contexts that remained stable.

**Figure S2.**
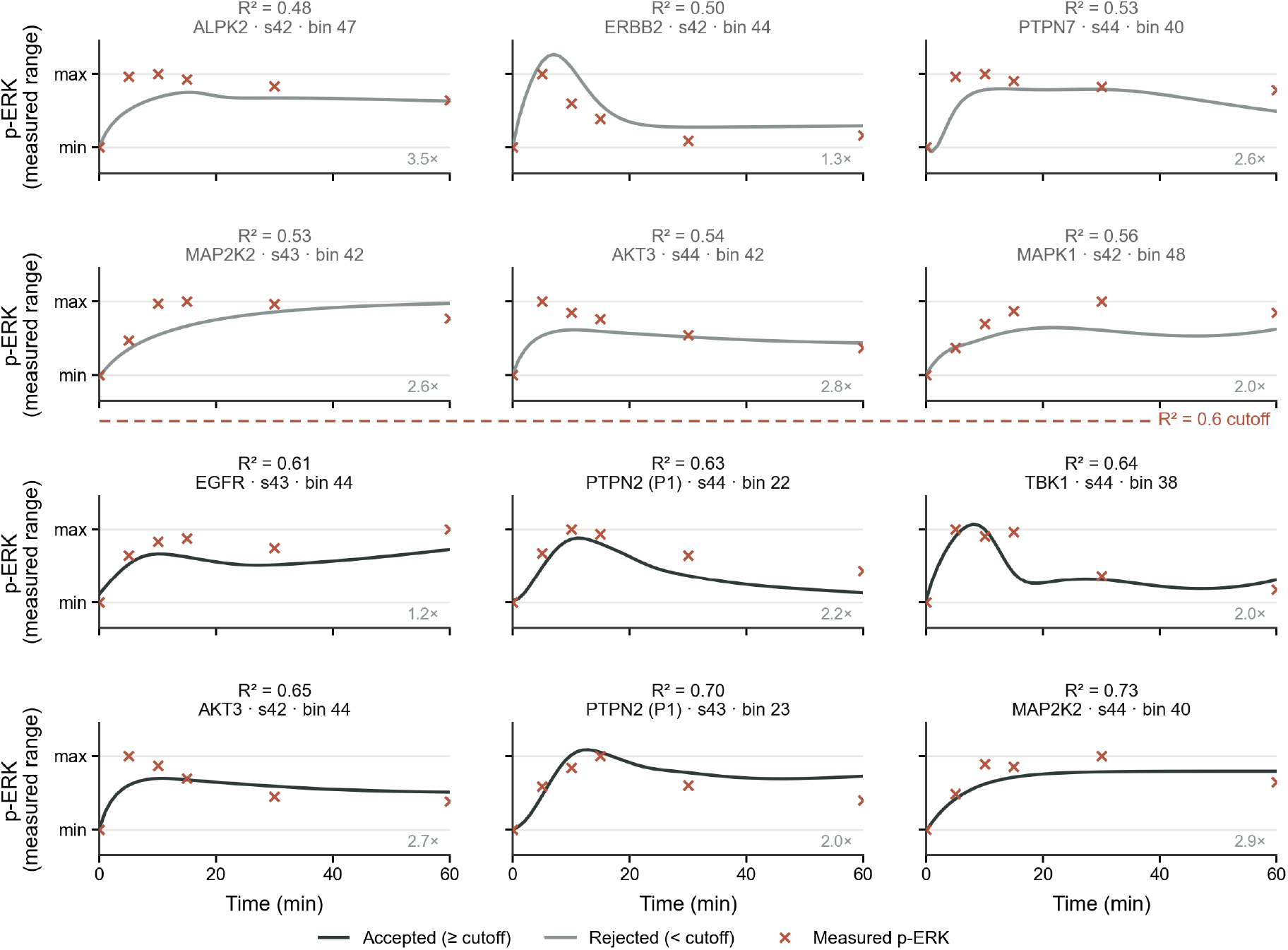
Representative integrated symbolic-regression trajectories near the R^2^ performance threshold. Fits are ranked by their test integrated R², the median across test GFP bins and the quantity the 0.6 cutoff is applied to; three fits are shown per target value (0.50, 0.55, 0.60, 0.65, 0.70), each at the test bin sitting at its median. Lines show the ODE-integrated symbolic-regression prediction (dark where the fit clears R² = 0.6, grey where it does not) and crosses the raw p-ERK measurements used for scoring. Trajectories at and above the cutoff reproduce the rise, peak timing and decay of the measured dynamics, whereas those below it flatten the peak or invert the decay. Generated under the selected configuration and the top-GFP-bin out-of-distribution split.

**Figure S3.**
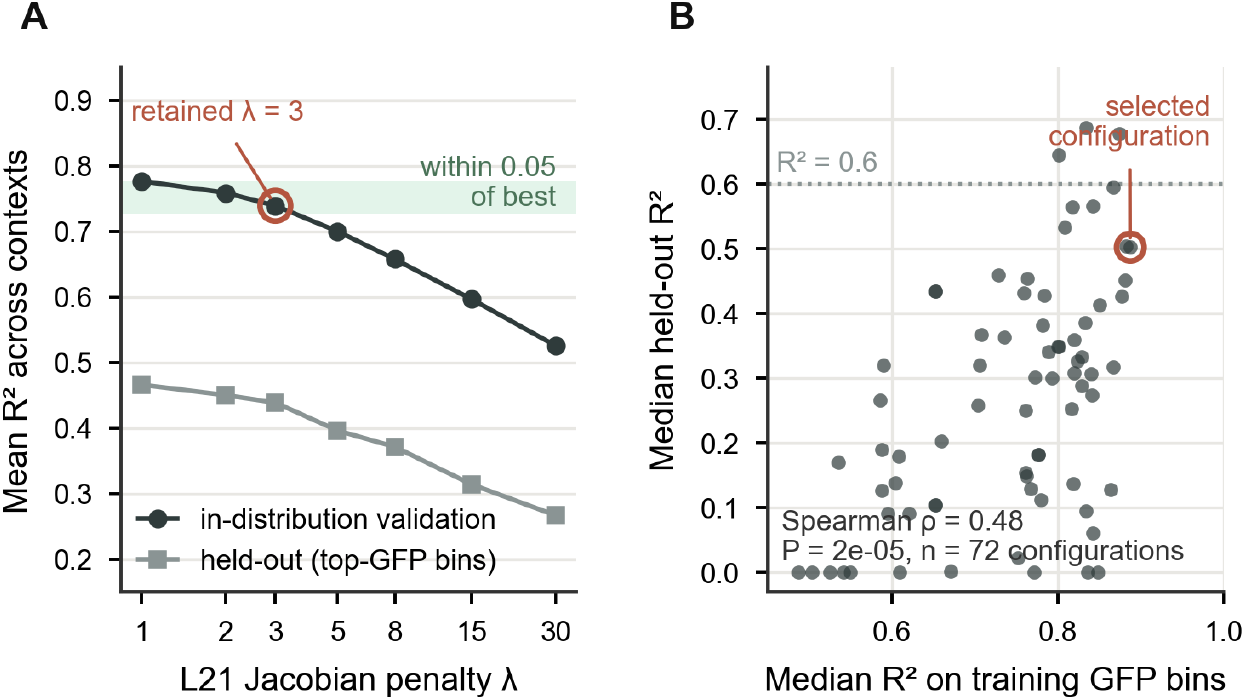
Sparse neural ODE regulariser and hyperparameter selection. (A) L21 λ sweep showing mean in-distribution validation R² and mean test R² across contexts. The elbow rule — the largest λ whose validation R² is within 0.05 of the best — retains λ = 3. (B) Median R² on the training GFP bins against median test R² across the 72-arm PySR configuration grid, the only grid here scored on both splits. In-distribution fit quality tracks test performance only loosely (Spearman ρ = 0.48, p=2×10^-5^), so the configuration selected on the training rule is not the best-extrapolating arm.

**Table S1.** Equations found by PySR across measurement noise degradation experiment (sQSSA lens)

| Regime | Seed 42 | Seed 43 | Seed 44 |
| --- | --- | --- | --- |
| 0% | $\frac{E_{tot}S}{\frac{k_{cat} + k_{inact} + 1.4 k_{off}}{K_D k_{off}} + S}$ | $\frac{E_{tot}S}{\frac{k_{off} + k_{cat} + k_{inact}}{K_D k_{off}} + S}$ | $\frac{0.26 E_{tot}S}{\frac{k_{off} + k_{cat} + k_{inact}}{3.8 K_D k_{off}} + S}$ |
| 1% | $\frac{0.11 E_{tot}S}{\frac{k_{cat} + k_{inact}}{9.0 K_D k_{off}} + S}$ | $\frac{E_{tot}S}{\frac{k_{cat} + k_{off} + k_{inact}}{K_D k_{off}} + S}$ | $\frac{E_{tot}S}{\frac{k_{inact} + k_{off} + k_{cat}}{K_D k_{off}} + S}$ |
| 10% | $\frac{E_{tot}S}{\frac{k_{inact} + 1.3 k_{off} + k_{cat}}{K_D k_{off}} + S}$ | $\frac{0.065 E_{tot}S}{\frac{k_{cat} + k_{inact}}{15 K_D k_{off}} + S}$ | $\frac{0.10 E_{tot}S}{\frac{k_{cat} + k_{inact}}{9.7 K_D k_{off}} + S}$ |
| 100% | $1.5 \times 10^{-4} E_{tot}$ | $1.5 \times 10^{-4} E_{tot}$ | $1.4 \times 10^{-4} E_{tot}$ |

**Table S2.** Equations found by PySR across dataset size degradation experiment (sQSSA lens)

| Regime | Seed 42 | Seed 43 | Seed 44 |
| --- | --- | --- | --- |
| 10% | $\frac{E_{tot}S}{\frac{1}{K_D} \left( \frac{k_{cat} + k_{inact}}{k_{off}} + 1.4 \right) + S}$ | $\frac{E_{tot} K_D k_{off} S}{k_{cat} + k_{off} + k_{inact}}$ | $\frac{E_{tot}S}{\frac{1}{K_D} \left( \frac{k_{inact} + k_{cat}}{k_{off}} + 1.3 \right) + S}$ |
| 50% | $\frac{0.086 E_{tot}S}{\frac{k_{cat} + k_{inact}}{12 K_D k_{off}} + S}$ | $\frac{E_{tot}S}{\frac{k_{cat} + k_{inact} + k_{off}}{K_D k_{off}} + S}$ | $\frac{E_{tot}S}{\frac{1}{K_D} \left( \frac{k_{cat} + k_{inact}}{k_{off}} + 1.3 \right) + S}$ |
| 100% | $\frac{0.77 E_{tot}S}{\frac{k_{cat} + k_{inact}}{1.3 K_D k_{off}} + S}$ | $\frac{0.079 E_{tot}S}{\frac{k_{inact} + k_{cat}}{13 K_D k_{off}} + S}$ | $\frac{E_{tot}S}{\frac{k_{off} + k_{inact} + k_{cat}}{K_D k_{off}} + S}$ |
| 200% | $\frac{E_{tot}S}{\frac{k_{cat} + k_{off} + k_{inact}}{K_D k_{off}} + S}$ | $\frac{E_{tot}S}{\frac{k_{inact} + k_{off} + k_{cat}}{K_D k_{off}} + S}$ | $\frac{E_{tot}S}{\frac{k_{inact} + k_{off} + k_{cat}}{K_D k_{off}} + S}$ |

**Table S3.** Equations found by PySR across Michaelis-Menten deviation degradation experiment (sQSSA lens)

| Regime | Seed 42 | Seed 43 | Seed 44 |
| --- | --- | --- | --- |
| $S/E_{tot} \gg 1$ | $\frac{E_{tot}S}{\frac{k_{cat} + k_{inact}}{K_D k_{off}} + S}$ | $\frac{0.16 E_{tot}S}{\frac{k_{cat} + k_{inact}}{6.3 K_D k_{off}} + S}$ | $\frac{0.16 E_{tot}S}{\frac{k_{inact} + k_{cat}}{6.1 K_D k_{off}} + S}$ |
| $S/E_{tot} \ll 1$ | $\frac{E_{tot} K_D k_{off} S}{k_{off} + k_{inact}}$ | $\frac{E_{tot} K_D S}{\frac{k_{inact}}{k_{off}} + 2.9}$ | $\frac{E_{tot} K_D k_{off} S}{k_{off} + k_{inact}}$ |
| $K_M/S \gg 1$ | $\frac{E_{tot} K_D k_{off} S}{k_{cat} + k_{inact} + k_{off}}$ | $\frac{E_{tot} K_D k_{off} S}{k_{cat} + k_{inact} + k_{off}}$ | $\frac{E_{tot} K_D k_{off} S}{k_{cat} + k_{inact} + k_{off}}$ |
| $K_M/S \ll 1$ | $\frac{97 E_{tot}}{\frac{k_{cat}}{k_{inact}} + 1.2 \times 10^2}$ | $\frac{E_{tot}}{k_{cat} + 1.5}$ | $\frac{57 E_{tot}}{\frac{k_{cat}}{k_{inact}} + 55}$ |

**Table S4.** Equations found by PySR across measurement noise degradation experiment (tQSSA lens)

| Regime | Seed 42 | Seed 43 | Seed 44 |
| --- | --- | --- | --- |
| 0% | $\frac{E_{tot}S}{\frac{k_{cat} + k_{inact}}{K_D k_{off}} + 16 S}$ | $E_{tot}S \sqrt{1.9 \times 10^{-6} K_D k_{off}}$ | $\frac{E_{tot}S}{\frac{k_{cat} + k_{inact}}{K_D k_{off}} + 13 \times 10^1}$ |
| 1% | $1.6 \times 10^{-3} K_D E_{tot} S$ | $\frac{E_{tot}S}{\frac{k_{cat} + k_{inact}}{K_D k_{off}} + 88}$ | $\frac{E_{tot}S}{\frac{k_{cat} + k_{inact}}{K_D k_{off}} + 10 \times 10^1}$ |
| 10% | $\frac{K_D E_{tot} S}{\frac{k_{cat} + k_{inact}}{k_{off}} + 34}$ | $\frac{K_D E_{tot} S}{K_D S + \frac{k_{cat} + k_{inact}}{k_{off}}}$ | $\frac{E_{tot}S}{\frac{k_{cat} + k_{inact}}{K_D k_{off}} + 35}$ |
| 100% | $8.5 \times 10^{-6} S$ | $5.5 \times 10^{-5} E_{tot} S$ | $6.4 \times 10^{-6} S$ |

**Table S5.** Equations found by PySR across dataset size degradation experiment (tQSSA lens)

| Regime | Seed 42 | Seed 43 | Seed 44 |
| --- | --- | --- | --- |
| 10% | $0.0020 K_D E_{tot} S$ | $0.0024 K_D E_{tot} S$ | $9.7 \times 10^{-5} E_{tot}$ |
| 50% | $0.0014 E_{tot} S \sqrt{K_D k_{off}}$ | $\frac{E_{tot} S}{(E_{tot} + S) + \frac{k_{inact} + k_{cat} + 1.3 k_{off}}{K_D k_{off}}}$ | $\frac{E_{tot} S}{E_{tot} + 3.7 S + \frac{0.92 (k_{inact} + k_{cat} + 1.3 k_{off})}{K_D k_{off}}}$ |
| 100% | $0.0014 K_D E_{tot} S$ | $\frac{E_{tot} S}{\frac{k_{inact} + k_{cat}}{K_D k_{off}} + 1.0 \times 10^2}$ | $\frac{E_{tot} S}{\frac{k_{inact} + k_{cat} + k_{off}}{K_D k_{off}} + (E_{tot} + S) + 0.016 S \frac{k_{cat}}{k_{inact}}}$ |
| 200% | $1.7 \times 10^{-4} E_{tot}$ | $0.0015 K_D E_{tot} S$ | $0.0025 E_{tot} S \sqrt{\frac{K_D k_{off}}{k_{cat} + k_{inact}}}$ |

**Table S6.**
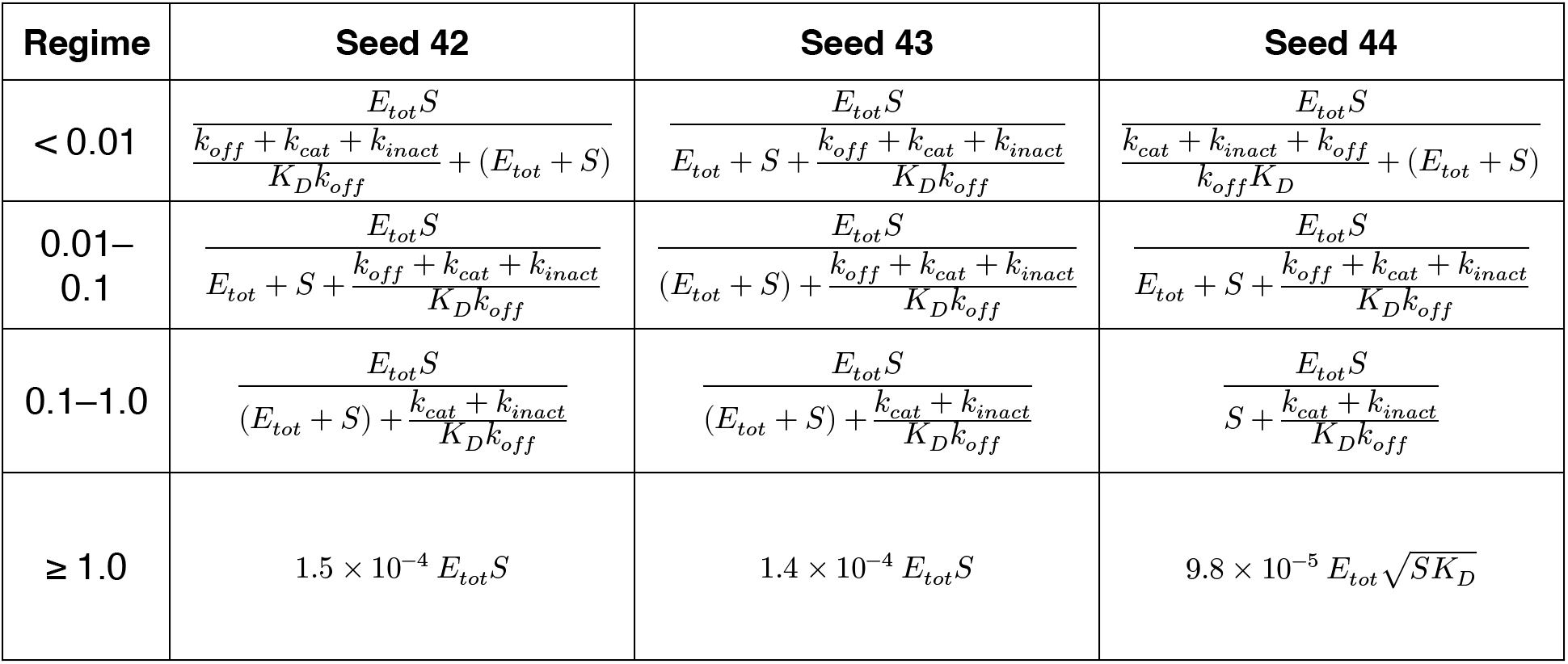
Equations found by PySR across Michaelis-Menten deviation degradation experiment (tQSSA lens)

**Table S7.**
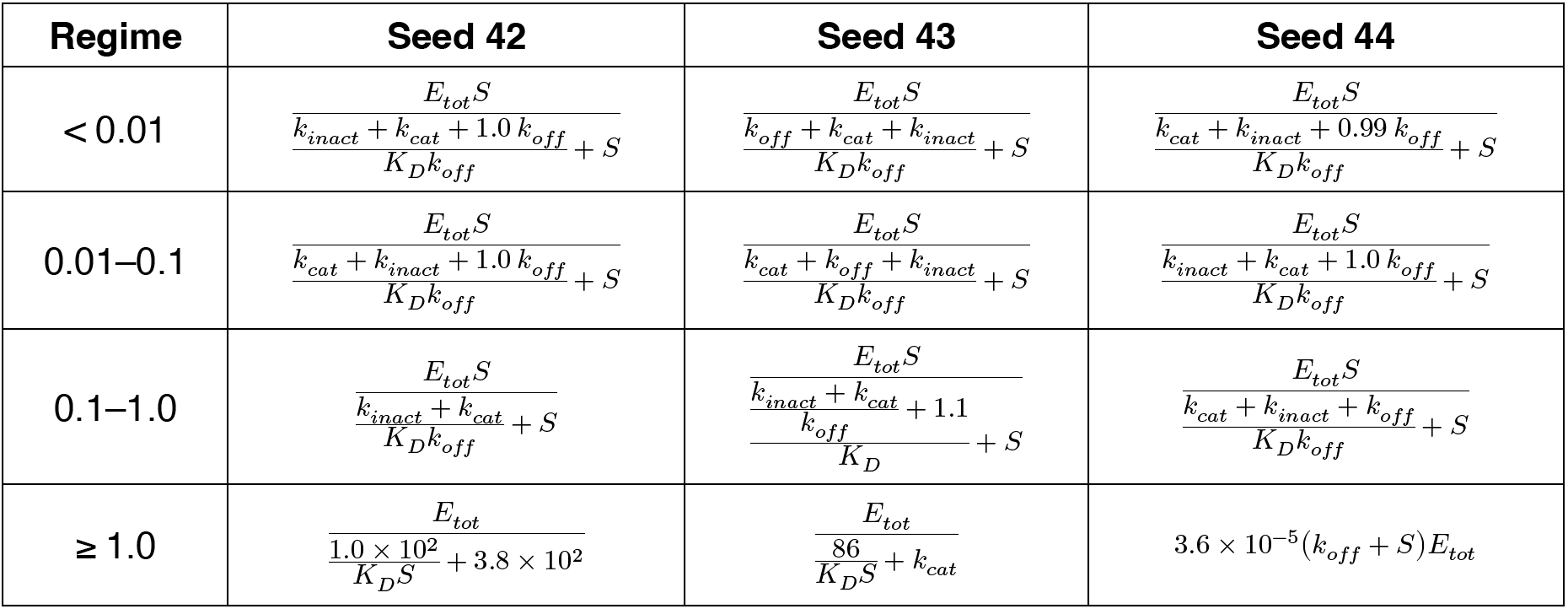
Equations found by PySR across kinetic limits degradation experiment (sQSSA lens)

**Table S8.** Symbolic-regression result per context. For each of the 40 contexts: the seed retained by training R², its training and test integrated R², whether it met the R² ≥ 0.6 criterion, and how many of the three seeds did. Variables used counts distinct model inputs appearing in the recovered law; complexity is its node count. The neural ODE’s test R², dependency count and interacting-pair count are shown alongside, the latter two averaged over three seeds. Rows are ordered by test R².

| Context | Seed | Train $R^2$ | Test $R^2$ | $\geq 0.6$ | Seeds $\geq 0.6$ | Variables | Complexity | NODE $R^2$ | NODE dep. | NODE pairs |
| --- | --- | --- | --- | --- | --- | --- | --- | --- | --- | --- |
| FLAG-GFP3 | 44 | 0.932 | 0.935 | yes | 3/3 | 5 | 18 | 0.948 | 3.67 | 5.33 |
| PIP5K3 | 44 | 0.932 | 0.928 | yes | 3/3 | 4 | 15 | 0.568 | 4.00 | 6.33 |
| untransfected3 | 42 | 0.957 | 0.924 | yes | 2/3 | 3 | 15 | 0.915 | 2.33 | 1.67 |
| untransfected2 | 44 | 0.948 | 0.915 | yes | 3/3 | 4 | 17 | 0.915 | 2.67 | 2.33 |
| PTPN7 | 43 | 0.906 | 0.910 | yes | 2/3 | 3 | 14 | 0.751 | 4.00 | 6.33 |
| FLAG-GFP2 | 43 | 0.896 | 0.801 | yes | 3/3 | 4 | 13 | 0.888 | 3.00 | 3.00 |
| untransfected1 | 42 | 0.948 | 0.794 | yes | 1/3 | 4 | 15 | 0.000 | 3.67 | 5.00 |
| ALPK2 | 43 | 0.879 | 0.729 | yes | 1/3 | 5 | 18 | 0.884 | 3.00 | 4.33 |
| MAP2K2 | 44 | 0.020 | 0.727 | yes | 1/3 | 2 | 8 | 0.613 | 3.00 | 3.33 |
| FLAG-GFP1 | 44 | 0.956 | 0.706 | yes | 1/3 | 5 | 16 | 0.691 | 3.00 | 3.67 |
| PTPN2 (P1) | 43 | 0.838 | 0.698 | yes | 2/3 | 4 | 17 | 0.000 | 3.67 | 5.00 |
| FLAG-GFP4 | 42 | 0.893 | 0.675 | yes | 3/3 | 6 | 16 | 0.843 | 3.33 | 4.00 |
| TBK1 | 44 | 0.644 | 0.636 | yes | 1/3 | 6 | 18 | 0.489 | 4.33 | 6.67 |
| ERBB2 | 44 | 0.969 | 0.363 | no | 0/3 | 3 | 10 | 0.380 | 3.67 | 4.67 |
| MAST2 | 42 | 0.845 | 0.246 | no | 0/3 | 5 | 16 | 0.000 | 4.33 | 7.33 |
| DYRK3 | 44 | 0.879 | 0.187 | no | 0/3 | 7 | 17 | 0.177 | 4.33 | 7.67 |
| MET | 42 | 0.044 | 0.099 | no | 0/3 | 4 | 15 | 0.727 | 5.00 | 9.67 |
| PRKACA | 43 | 0.612 | 0.095 | no | 0/3 | 4 | 13 | 0.788 | 4.67 | 11.67 |
| ABL1 | 42 | 0.871 | 0.000 | no | 0/3 | 4 | 13 | 0.362 | 4.33 | 6.33 |
| AKT3 | 43 | 0.924 | 0.000 | no | 1/3 | 6 | 20 | 0.000 | 3.67 | 5.00 |
| ARAF | 44 | 0.749 | 0.000 | no | 0/3 | 4 | 16 | 0.000 | 3.00 | 3.00 |
| DUSP10 (P2) | 42 | 0.924 | 0.000 | no | 0/3 | 5 | 16 | 0.000 | 4.67 | 8.33 |
| DUSP16 | 42 | 0.603 | 0.000 | no | 0/3 | 3 | 12 | 0.887 | 3.67 | 5.00 |
| DUSP4 | 43 | 0.718 | 0.000 | no | 0/3 | 4 | 14 | 0.000 | 4.67 | 8.33 |
| DUSP7 | 42 | 0.000 | 0.000 | no | 0/3 | 2 | 6 | 0.771 | 4.00 | 6.00 |
| DYRK2 | 42 | 0.208 | 0.000 | no | 0/3 | 6 | 17 | 0.224 | 4.33 | 7.33 |
| EGFR | 42 | 0.000 | 0.000 | no | 1/3 | 5 | 20 | 0.889 | 3.00 | 3.33 |
| FGFR1 | 42 | 0.000 | 0.000 | no | 0/3 | 2 | 7 | 0.000 | 5.33 | 10.67 |
| MAP4K2 | 43 | 0.275 | 0.000 | no | 0/3 | 5 | 18 | 0.000 | 3.33 | 4.33 |
| MAPK1 | 44 | 0.400 | 0.000 | no | 0/3 | 7 | 16 | 0.970 | 5.00 | 10.00 |
| MAPK3 | 42 | 0.620 | 0.000 | no | 0/3 | 5 | 14 | 0.925 | 5.33 | 9.00 |
| MST1R | 44 | 0.504 | 0.000 | no | 0/3 | 5 | 16 | 0.278 | 4.00 | 5.67 |
| PIK3R1 | 42 | 0.970 | 0.000 | no | 0/3 | 5 | 16 | 0.000 | 3.67 | 5.33 |
| PTPN5 | 42 | 0.560 | 0.000 | no | 0/3 | 5 | 19 | 0.000 | 4.00 | 5.67 |
| RPS6KA1 | 42 | 0.000 | 0.000 | no | 0/3 | 4 | 12 | 0.000 | 4.00 | 6.00 |
| RPS6KA3 | 42 | 0.000 | 0.000 | no | 0/3 | 4 | 10 | 0.000 | 2.67 | 2.67 |
| RPS6KA6 | 42 | 0.000 | 0.000 | no | 0/3 | 2 | 6 | 0.478 | 3.00 | 3.00 |
| TEC | 43 | 0.688 | 0.000 | no | 0/3 | 5 | 21 | 0.000 | 5.33 | 12.67 |
| TYRO3 | 42 | 0.000 | 0.000 | no | 0/3 | 5 | 16 | 0.347 | 5.00 | 11.00 |
| untransfected4 | 43 | 0.873 | 0.000 | no | 0/3 | 5 | 22 | 0.882 | 2.33 | 1.67 |

**Table S9.** PySR configuration sweep. Seventy-two configurations (max-iterations ∈ {700, 1400} × populations ∈ {30, 60} × population size ∈ {30, 60} × maximum complexity ∈ {18, 22, 26} × parsimony ∈ {0.1, 0.8, 3.0}), binary operators fixed to {+, −, ×, ÷} and no unary operators, each run on the six-context development set with three seeds (18 fits per configuration, 1,296 in total). Each row gives the median across the arms sharing that hyperparameter value; the last column flags the values of the selected configuration. Configurations were ranked by median R² on the training GFP bins, a rule fixed before results were seen, so the test columns played no part in the choice—they are shown to make clear that the two agree on maximum complexity 26 and parsimony 0.8 but not on population count. The full per-configuration grid is deposited with the code.

| Hyperparameter | Value | Arms | Median train $R^2$ | Median test $R^2$ | Contexts $\geq 0.6$ | Selected |
| --- | --- | --- | --- | --- | --- | --- |
| Max iterations | 700 | 36 | 0.764 | 0.220 | 6 |  |
| Max iterations | 1400 | 36 | 0.795 | 0.305 | 6 | yes |
| Populations | 30 | 36 | 0.721 | 0.186 | 6 | yes |
| Populations | 60 | 36 | 0.801 | 0.345 | 6 |  |
| Population size | 30 | 36 | 0.776 | 0.182 | 6 | yes |
| Population size | 60 | 36 | 0.778 | 0.327 | 6 |  |
| Max complexity | 18 | 24 | 0.733 | 0.146 | 6 |  |
| Max complexity | 22 | 24 | 0.776 | 0.287 | 6 |  |
| Max complexity | 26 | 24 | 0.819 | 0.349 | 7 | yes |
| Parsimony | 0.1 | 24 | 0.821 | 0.235 | 6 |  |
| Parsimony | 0.8 | 24 | 0.817 | 0.330 | 6 | yes |
| Parsimony | 3 | 24 | 0.652 | 0.172 | 6 |  |

**Table S10.**
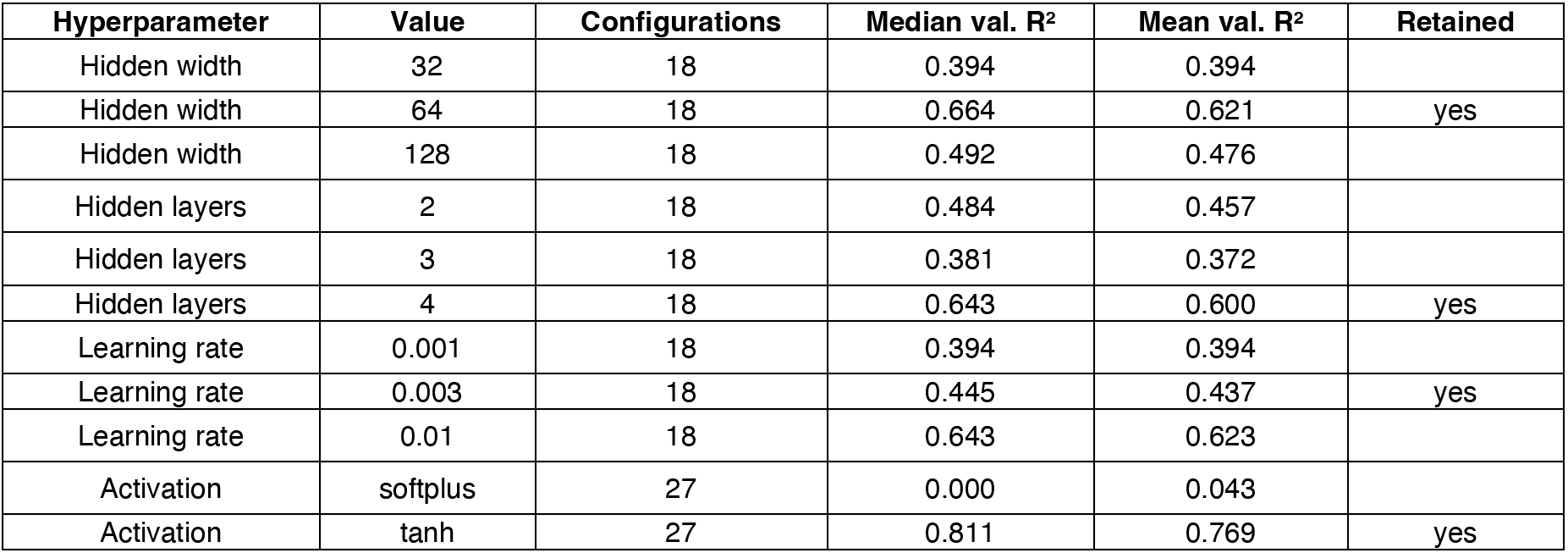
Sparse neural ODE architecture sweep. Fifty-four configurations (hidden width ∈ {32, 64, 128} × hidden layers ∈ {2, 3, 4} × learning rate ∈ {10^-3^, 3×10^-3^, 10^-2^} × activation ∈ {tanh, softplus}) were trained without sparsity regularisation and scored by in-distribution validation R² with a single seed. Each row gives the median and mean validation R² across the configurations sharing that hyperparameter value, so the table reads as the marginal effect of each choice; the last column flags the values of the retained configuration (width 64, 4 layers, 3×10^-3^, tanh). The retained learning rate is not the strongest marginal value (3×10^-3^ against 10^-2^) because the configuration was chosen on the best single arm rather than on these marginals.

**Table S11.**
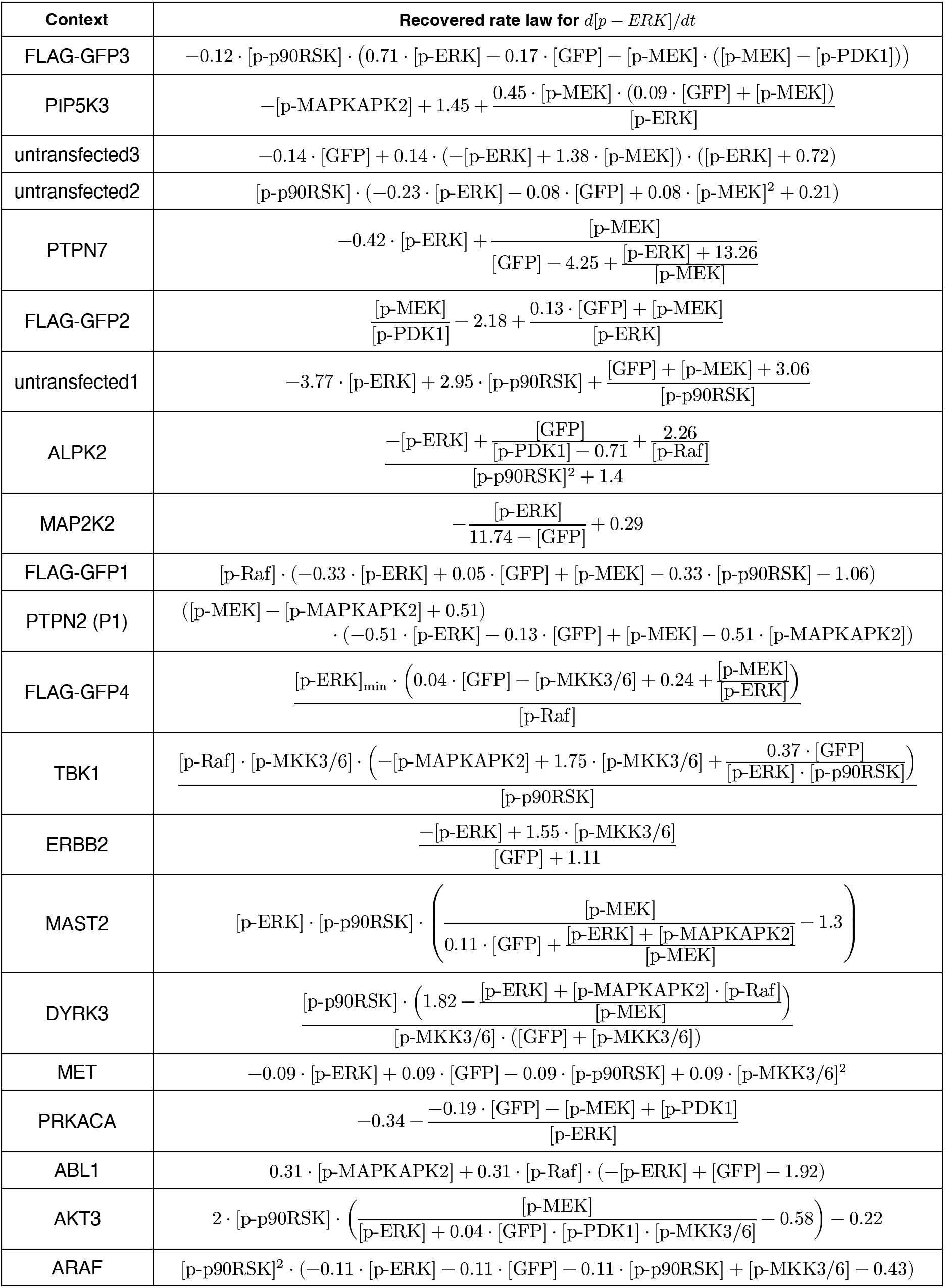
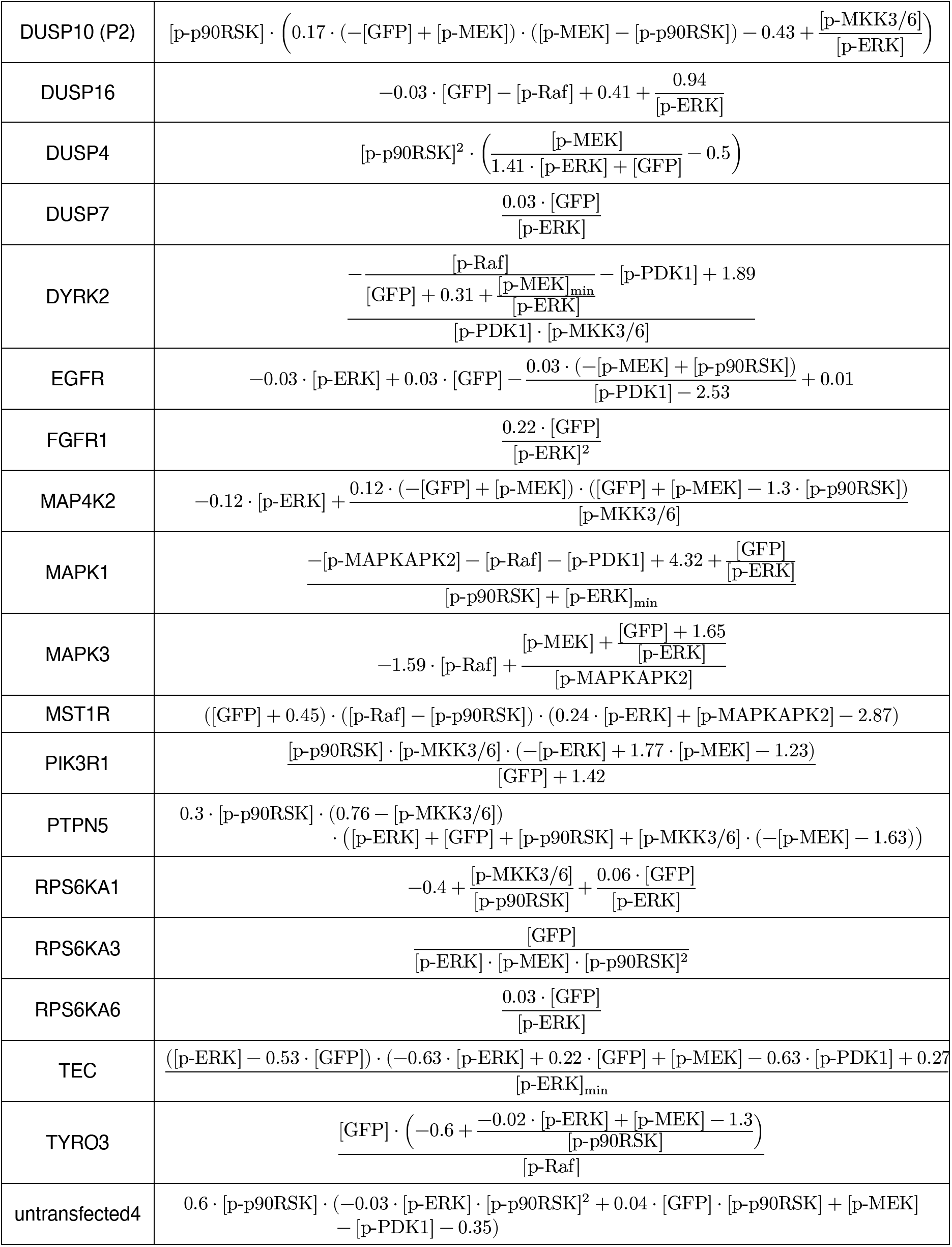
Recovered models by SR. The expression retained for each of the 40 contexts—the seed with the highest training integrated R², the same seed listed in Table S8—written for d[p-ERK]/dt. Square-bracketed names are the ten model inputs; subscript min denotes the baseline offset along the reconstructed trajectory. Coefficients are rounded to two decimal places, so re-integrating these expressions reproduces the reported trajectories only approximately.

**Table S12.** Sensitivity of the dependency-count contrast to the Jacobian threshold. Effective dependencies are inputs whose mean absolute Jacobian exceeds a fixed fraction of the largest; the analysis uses 0.15 throughout. Columns give the mean dependency count of contexts where symbolic regression failed and where it succeeded, with the accuracy-residualised permutation P among the 17 contexts the neural ODE fits, and the two-sided Mann–Whitney P across all 40. The contrast is significant from 0.05 to 0.25 on the gated comparison and from 0.05 to 0.20 across all 40; at 0.30 nearly every context collapses to about two inputs and the comparison loses its resolution.

| Jacobian threshold | Gated (17 contexts) |  |  | All 40 contexts |  |  |
| --- | --- | --- | --- | --- | --- | --- |
|  | Failed | Solved | P | Failed | Solved | P |
| 0.05 | 7.33 | 5.96 | 0.0107 | 7.57 | 6.51 | 0.0136 |
| 0.1 | 5.00 | 4.11 | 0.0462 | 5.12 | 4.41 | 0.0338 |
| 0.15 | 4.12 | 3.11 | 0.0248 | 4.09 | 3.36 | 0.0091 |
| 0.2 | 3.42 | 2.56 | 0.0251 | 3.33 | 2.82 | 0.0399 |
| 0.25 | 3.12 | 2.30 | 0.0225 | 2.79 | 2.44 | 0.1312 |
| 0.3 | 2.54 | 2.11 | 0.2936 | 2.25 | 2.23 | 0.9415 |

**Table S13.** Sensitivity of the interacting-pair contrast to the off-diagonal Hessian threshold. Interacting pairs are off-diagonal input-Hessian entries exceeding a fixed fraction of the largest such entry, counted among the inputs the Jacobian criterion retains; the Jacobian cut is held at 0.15 throughout so the two thresholds are not confounded, and the analysis uses 0.25. Ratio is failed over solved among the 17 contexts the neural ODE fits, with the accuracy-residualised permutation P; the last columns give the two-sided Mann–Whitney P across all 40. Failed contexts carry more interacting pairs at every threshold tested, significantly so from 0.05 to 0.30, with the ratio declining as the cut rises and fewer pairs survive in either group.

| Off-diagonal threshold | Gated (17 contexts) |  |  |  | All 40 contexts |  |  |
| --- | --- | --- | --- | --- | --- | --- | --- |
|  | Failed | Solved | Ratio | P | Failed | Solved | P |
| 0.05 | 7.58 | 3.81 | 1.99× | 0.0212 | 7.33 | 4.56 | 0.0144 |
| 0.1 | 7.58 | 3.81 | 1.99× | 0.0212 | 7.33 | 4.54 | 0.0123 |
| 0.15 | 7.46 | 3.81 | 1.96× | 0.0210 | 7.25 | 4.54 | 0.0151 |
| 0.2 | 7.37 | 3.81 | 1.93× | 0.0209 | 7.14 | 4.51 | 0.0157 |
| 0.25 | 7.04 | 3.78 | 1.86× | 0.0231 | 6.72 | 4.38 | 0.0192 |
| 0.3 | 6.29 | 3.67 | 1.72× | 0.0329 | 6.06 | 4.23 | 0.0270 |
| 0.4 | 5.38 | 3.33 | 1.61× | 0.0561 | 5.11 | 3.74 | 0.0544 |
| 0.5 | 4.17 | 2.78 | 1.50× | 0.0807 | 4.12 | 3.03 | 0.0555 |

**Table S14.** Calibration of the L21 penalty weight. Each λ was trained across 40 contexts and three seeds, and scored by mean in-distribution validation R² — the quantity the selection rule acts on — with test R² shown alongside for reference only. The elbow rule retains the largest λ whose mean validation R² falls within 0.05 of the best, giving λ = 3. Because a stronger penalty yields a sparser network, this selects the sparsest model that is not measurably worse, so the dependency counts reported throughout are conservative with respect to sparsity: a weaker penalty would raise them. Data are deposited as table_s14_lambda_calibration.csv.

| $\lambda$ | Mean val. $R^2$ | Median val. $R^2$ | Mean test $R^2$ | Median test $R^2$ | Within 0.05 of best | Retained |
| --- | --- | --- | --- | --- | --- | --- |
| 1 | 0.777 | 0.881 | 0.466 | 0.404 | yes | no |
| 2 | 0.759 | 0.846 | 0.450 | 0.449 | yes | no |
| 3 | 0.739 | 0.836 | 0.439 | 0.430 | yes | yes |
| 5 | 0.700 | 0.798 | 0.396 | 0.416 | no | no |
| 8 | 0.658 | 0.729 | 0.371 | 0.334 | no | no |
| 15 | 0.597 | 0.655 | 0.314 | 0.233 | no | no |
| 30 | 0.525 | 0.586 | 0.267 | 0.191 | no | no |

